# GOT1 primes the cellular response to hypoxia by supporting glycolysis and HIF1α stabilisation

**DOI:** 10.1101/2023.09.06.556522

**Authors:** Fiona Grimm, Agustín Asuaje, Aakriti Jain, Mariana Silva dos Santos, Jens Kleinjung, Patrícia M. Nunes, Stefanie Gehrig, Louise Fets, James I. MacRae, Dimitrios Anastasiou

**Affiliations:** Cancer Metabolism Laboratory, The Francis Crick Institute, 1 Midland Road, NW1 1AT, London, United Kingdom; Metabolomics Science Technology Platform, The Francis Crick Institute, 1 Midland Road, NW1 1AT, London, United Kingdom; Computational Biology Science Technology Platform, The Francis Crick Institute, 1 Midland Road, NW1 1AT, London, United Kingdom

**Keywords:** hypoxia, metabolism, glycolysis, HIF1α, GOT1, α-ketoglutarate

## Abstract

Adaptation to chronic hypoxia occurs through changes in protein expression, which are controlled by hypoxia inducible factor 1a (HIF1α) and are necessary for cancer cell survival. However, the mechanisms that enable cancer cells to adapt in early hypoxia, prior to full activation of HIF1α, remain poorly understood. Here we show that aspartate transaminase 1 (GOT1), which supports NAD^+^ production by malate dehydrogenase 1 (MDH1), is required, in addition to reserve lactate dehydrogenase (LDH) capacity, for the HIF1α-independent increase in glycolysis we observe early upon exposure of cells to hypoxia. Additionally, GOT1 maintains low α-ketoglutarate levels, thereby limiting prolyl hydroxylase activity to promote HIF1α stabilisation in early hypoxia and robust HIF1α target gene expression in later hypoxia. Our findings reveal that, in normoxia, GOT1 maintains cells in a primed state and ready to support increased glycolysis and HIF1α stabilisation upon oxygen limitation, until other adaptive processes that require more time, are fully established.

## INTRODUCTION

Glucose metabolism is important for many physiological and pathological processes. In cancer, glycolysis is often increased though multiple mechanisms that include upregulated expression of glucose transporters and glycolytic genes, differential expression of metabolic enzyme isoforms, and aberrant oncogenic signalling ^1–3^. These mechanisms promote both glucose transport into cells as well as increased glycolytic enzyme activity, which, collectively, enhance glucose metabolism ^4^. Furthermore, a high oxidised-to-reduced nicotinamide adenine dinucleotide (NAD^+^/NADH) ratio is also required to sustain the NAD^+^-dependent activity of the glycolytic enzyme glyceraldehyde 3-phosphate dehydrogenase (GAPDH), which becomes rate-limiting when glycolysis is high ^5,6^.

The function of increased glycolysis in tumours remains under intense investigation. Although glucose metabolism can provide precursors for biosynthetic pathways, a relatively low proportion of glucose carbons enters biomass production ^7^. However, there is significant evidence that a major role of glycolysis is to maintain energy balance by producing ATP under conditions that perturb cellular bioenergetics ^8–10^. Accordingly, increased glucose uptake correlates well with hypoxic tumour regions ^11–13^, where ATP production from mitochondria is attenuated due to limiting oxygen ^14,15^.

During chronic hypoxia, upregulation of glycolysis is achieved through a co-ordinate increase in glycolytic enzyme expression ^16^, which is orchestrated by the prolyl hydroxylase (PHD)-hypoxia inducible factor 1α (HIF1α) signalling axis ^17–19^. PHDs use iron, α-ketoglutarate (αKG) and O_2_ to hydroxylate proline residues on HIF1α, which are then recognised by the E3 ligase von Hippel Lindau (VHL), leading to its ubiquitination and subsequent degradation by the proteasome ^20–25^. The K_m_ of PHDs for O_2_ lies within the physiologically relevant range of O_2_ concentrations in tissues, therefore these enzymes are thought to function as oxygen sensors ^26–30^. Binding of O_2_ to the catalytic pocket of PHDs requires prior binding of αKG ^25^, which also prevents re-association of hydroxylated HIF1α to PHDs, enabling more efficient HIF1α degradation ^31^. Some oncometabolites can outcompete αKG binding to PHDs, leading to HIF stabilisation ^32–35^, and this can be alleviated by exogenous αKG ^36–38^. Consequently, in addition to fluctuations in O_2_ and post-translational modification of PHD catalytic residues ^39,40^, αKG levels may also determine the turnover kinetics of HIF1α. However, it is not known which of the pathways involved in αKG metabolism regulate HIF1α expression dynamics during the onset of hypoxia.

Upon its stabilisation in hypoxia, HIF1α controls the transcription of genes that include glucose transporters and most glycolytic genes ^41^. Concomitantly, HIF1α drives the expression of pyruvate dehydrogenase kinase 1 (PDK1), which catalyses the inhibitory phosphorylation of pyruvate dehydrogenase (PDH), leading to decreased entry of glucose carbons into the tricarboxylic acid (TCA) cycle ^42,43^. It has been postulated that decreased TCA cycle activity attenuates mitochondrial NAD^+^-regenerating pathways, such as the malate-aspartate shuttle (MAS), leading to increased reliance of glycolysis on lactate dehydrogenase A (LDHA) for NAD^+^ ^18,44^. Increased availability of pyruvate, the LDHA substrate, in the cytoplasm following PDH inhibition promotes LDHA activity ^45^. Moreover, the *LDHA* gene is also a HIF1α target, resulting in enhanced LDHA protein expression in hypoxia to further increase NAD^+^ production _46_. Accordingly, cells that rely more on glycolysis are more sensitive to inhibition of LDHA compared to cells that depend on mitochondria for ATP production ^47^. Furthermore, knock-down or pharmacological inhibition of LDHA in hypoxic cancer cells results in decreased proliferation and leads to cell death attributed to oxidative stress ^48–51^. Collectively, this evidence indicates that, increased LDHA protein expression, in addition to that of glucose transporters and glycolytic enzymes, is also required for increased glycolysis in chronic hypoxia ^52,53^.

Intriguingly, while changes in gene expression through HIF1α, or other mechanisms, require more than 24 h to reach maximal levels ^54^, upregulation of glycolysis occurs within minutes to hours upon exposure to hypoxia and is essential for sustaining ATP levels under these conditions ^15,55–57^. Acute increase in glycolysis upon hypoxia, or after inhibition of mitochondrial respiration, is due to the reversal of the Pasteur effect, which describes the inhibitory effect of oxygen on glycolysis. The Pasteur effect is mediated, in part, by increased activity of phosphofructokinase (PFK) due to decreased production of its allosteric inhibitor ATP in mitochondria ^58,59^, and increased phosphorylation by adenosine monophosphate-activated protein kinase (AMPK) ^60,61^. Moreover, oxygen limitation can also upregulate glycolysis by directly influencing glucose uptake through mechanisms that include modification of glucose transporters or their increased translocation to the plasma membrane ^55,56,62–65^. Although our understanding of the processes involved in the initiation of the Pasteur effect is substantial, the mechanisms used to provide enough NAD^+^ to support the upregulation of glycolysis during the onset of hypoxia remain elusive. In particular, in light of the apparent need for increased expression of LDHA in chronic hypoxia, it is unclear whether basal LDHA expression suffices to sustain redox balance also in early hypoxia, prior to HIF1α-mediated effects, or whether other mechanisms exist to support the cellular requirements for NAD^+^ upon acute oxygen limitation.

Here we show that GOT1 (glutamate-oxoglutarate transaminase 1, also known as aspartate aminotransferase 1) supports malate dehydrogenase 1 (MDH1), which becomes an important source of NAD^+^ during the first hours of hypoxia, when glycolysis increases and reserve LDHA does not suffice to sustain maximal GAPDH activity. Additionally, GOT1 consumes αKG, leading to attenuated PHD activity and increased HIF1α stabilisation in early hypoxia, and robust HIF1α target gene expression in later hypoxia. Our findings reveal a role for GOT1 activity in priming cells for the first hours in hypoxia and until other adaptive mechanisms involved in chronic hypoxia are fully established.

## RESULTS

### Glycolysis increases within 3 h in hypoxia and correlates with decreased aspartate levels

To investigate metabolic changes elicited by low oxygen concentrations, we measured intracellular metabolites in MCF7 cells incubated in 21% (normoxia) or 1% O_2_ (hypoxia) for increasing lengths of time, between 1-24 h (**Figure 1A**). The earliest and most statistically significant changes we observed were an increase in lactate and a decrease in intracellular aspartate levels, both of which persisted into later time points (**Figure 1B-D**). These changes also occurred, to varying degrees, in other breast cancer cell lines, as well as immortalised, non-tumorigenic mammary epithelial cells (**Figure S1A-B**). Upon re-oxygenation following 3 h in hypoxia, lactate and aspartate levels in MCF7 cells returned to pre-hypoxia levels with comparable, albeit slower, kinetics than their onset, indicating that hypoxia-induced changes in lactate and aspartate are reversible (**Figure S1C-D**). Treatment of cells with antioxidants did not attenuate the increase in lactate or decrease in aspartate (**Figure S1E-G**), negating an involvement of increased reactive oxygen species (ROS), which have other important signalling roles in early hypoxia ^66,67^. Accumulation of intracellular lactate coincided with increased cellular glucose uptake at 1% O_2_ [(1.69 ± 0.02)-fold and (2.78 ± 0.78)-fold after 3 h and 24 h, respectively] (**Figure 1E**) and was accompanied by elevated lactate excretion into the media (**Figure 1F**). Isotopic labelling with [U-^13^C]-glucose showed that ^13^C-labelling of secreted lactate was higher in hypoxia (**Figure S1H**) indicating that increased lactate is due to enhanced glycolysis. Together, these data showed that increased glycolysis occurs within 3 h after cells are exposed to 1% O_2_ and coincides with a decrease in aspartate levels.

**Figure 1.**
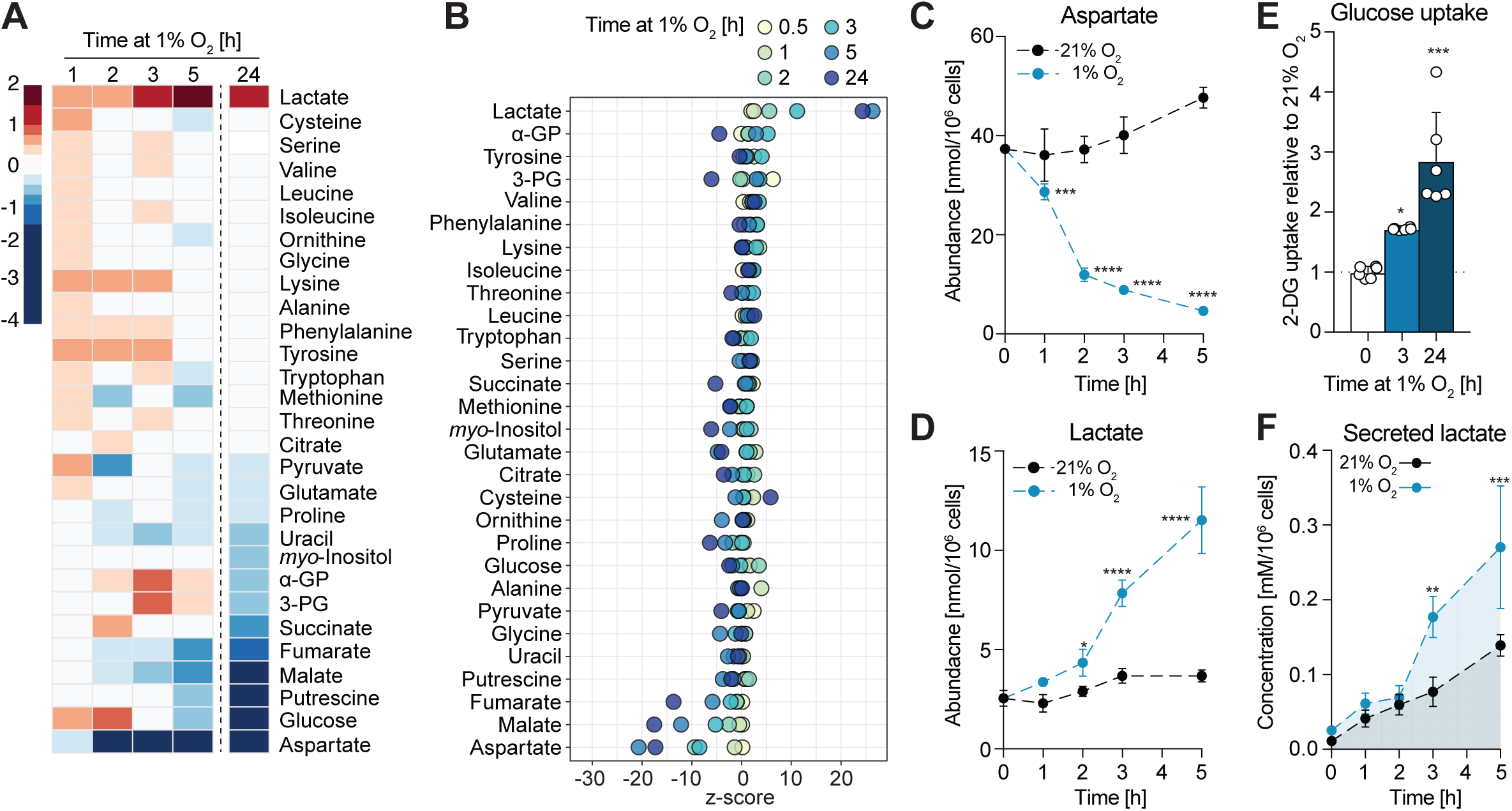
Increased glycolysis occurs within 3 h upon exposure to 1% O_2_ and correlates with decreased intracellular aspartate levels. **A**) Heatmap showing log_2_ fold-changes of the abundance of the indicated metabolites in MCF7 cells exposed to 1% O_2_ for the indicated lengths of time, compared to cells at 21% O_2_ (n = 4 cultures per time point and condition). Metabolites are ordered according to log_2_ fold-changes after 24 h in 1% O_2_. B) Z score plot of changes in metabolite abundances shown in **A**. **C-D)** Intracellular abundances of lactate and aspartate shown in **A**. Data are shown as mean ±SD. Significance relative to 21% O_2_ was tested using two-way ANOVA with Sidak’s multiple comparison test. See also **Figure S1A-B**. **E)** Glucose (2DG) uptake of MCF7 cells in 21% O_2_ and after 3 or 24 h in 1% O_2_. Data are shown as mean ±SD (n = 6 assays per condition). Significance relative to 21% O_2_ was tested using one-way ANOVA with Dunnett’s multiple comparisons test. **F)** Lactate concentration in culture media of MCF7 cells incubated in 21% O_2_ or 1% O_2_ for the indicated lengths of time (n = 4 cultures per time point and condition). Data are shown as mean ±SD and significance relative to 21% O_2_ was tested using two-way ANOVA with Sidak’s multiple comparison test. Significance levels in all figures are: * p<0.05, ** p<0.01, *** p<0.001, **** p<0.0001.

### Early increase in glycolysis is not dependent on HIF1α

Upregulation of glycolysis in chronic hypoxia is commonly attributed to the transcriptional activity of HIF1α, which results in increased glucose uptake and lactate production ^68^. We found that HIF1α protein levels increased within 1 h and reached maximal levels within 3 h in hypoxia in all cell lines tested (**Figure 2A** left, **Figure S2A**). Expression of HIF2α, which also has important roles in the regulation of gene expression in hypoxia ^69^, did not change detectably within the time-frame tested. Although changes in mRNA expression of many known HIF1α target genes ^70^ were detected in MCF7 cells after 3 h at 1% O_2_, transcriptional up- and down-regulation of most genes within this panel was more pronounced after 24 h in hypoxia (**Figure 2B**). Despite the early onset of the transcriptional response (within 3 h in hypoxia), changes in protein levels of HIF1α targets involved in glucose uptake (GLUT1), glycolysis (HK2, PKM2) and lactate production (LDHA) were only detected after 6 h, but not after 3 h in hypoxia (**Figure 2A** left). Therefore, the early metabolic changes described above occurred before robust expression of HIF1α target genes involved in glycolysis and lactate production was detectable on the protein level.

**Figure 2.**
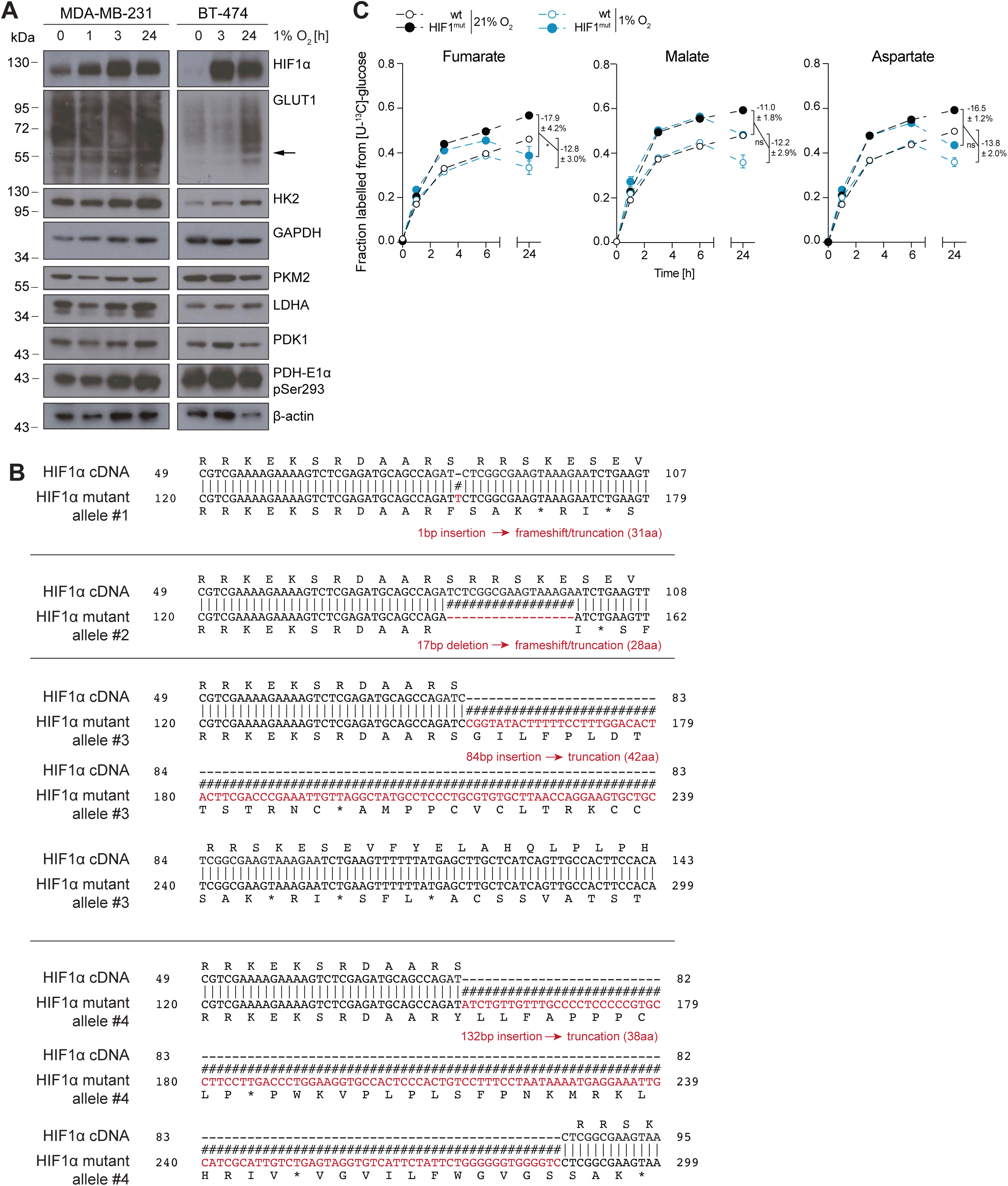
Increased glycolysis and depletion of aspartate in early hypoxia are independent of HIF1α. **A)** Western blot to assess levels of HIF1α and a panel of HIF1α targets in wild-type (wt) MCF7 and HIF1α^mut^ MCF7 cells incubated in 21% O_2_ or at 1% O_2_ for the indicated lengths of time. The asterisk marks a HIF1α immuno-reactive band of smaller molecular weight than HIF1α that increases upon hypoxia in HIF1α^mut^ MCF7 cells, indicative of a truncated HIF1α that likely lacks the transactivation domain where the sgRNA sequence is targeted at. See also **Figure S2A**. B) Log_2_ fold-changes in mRNA expression levels of a panel of HIF1α targets in MCF7 cells exposed to 1% O_2_ for 3 or 24 h, compared to control cells at 21% O_2_ (n = 3 cultures for each time point and condition; only changes with FDR<0.01 are shown). C) Heatmap showing log_2_ fold-changes in mRNA expression levels of a panel of HIF1α targets in wild-type (wt) and HIF1α^mut^ MCF7 cells exposed to 1% O_2_ for 3 or 24 h, compared to control cells in normoxia (n = 3 cultures for each time point and condition – only changes with FDR<0.01 are shown). D) Fraction of citrate labelled from [U-^13^C]-glucose in wild-type (wt) and HIF1α^mut^ MCF7 cells after incubation with the tracer at 21% O_2_ or 1% O_2_ for the indicated lengths of time. Time points indicate both the duration of hypoxia treatment and incubation with the tracer (n = 4 cultures for each time point and condition). Data are shown as mean ± SD. Statistical errors were propagated to calculate variance of the change in isotopic labelling between normoxia and hypoxia for each cell line. Significance of this change between cell lines was then tested using two-way ANOVA with Sidak’s multiple comparison test. See also **Figure S2C**. E) Changes in lactate and aspartate abundance in wild-type (wt) and HIF1α^mut^ MCF7 cells incubated in 21% O_2_ or 1% O_2_ for the indicated lengths of time, compared to control cells in normoxia (n = 4 cultures for each time point and condition). Data are shown as mean ± SD and significance relative to wild-type cells was tested using two-way ANOVA with Sidak’s multiple comparison test.

To confirm that the early upregulation of glycolysis was independent of HIF1α transcriptional activity, we engineered MCF7 cells that lack functional HIF1α (henceforth referred to as HIF1α^mut^ cells) using CRISPR/Cas9 (**Figure S2B**). HIF1α^mut^ cells showed a severe impairment in upregulating HIF1α target genes in hypoxia, both on the mRNA and protein level (**Figure 2A** right, **Figure 2C**). Moreover, decreased entry of glucose carbons into the TCA cycle, which occurs in a HIF1α-dependent manner in chronic hypoxia ^42^, was partially attenuated in HIF1α^mut^ cells after 24 h in 1% O_2_ (**Figure 2D, Figure S2C**). Similarly, lactate accumulation at later time points (6 and 24 h in 1% O_2_), was partially suppressed in HIF1α^mut^cells compared to wild-type MCF7 cells (**Figure 2E**). However, within the first 3 h in hypoxia, HIF1α^mut^ cells showed similar increases in lactate and decreases in aspartate as wild-type MCF7 cells (**Figure 2E**). Together, these data demonstrated that, while later metabolic changes are, at least partially, dependent on HIF1α, the early increase in glycolysis upon hypoxia treatment occurs independently of HIF1α transcriptional activity. Henceforth, we refer to 3 h hypoxia as “early hypoxia” to distinguish it from other, previously described, acute responses mediated by ROS ^66,67^.

### Knock-out of GOT1 attenuates the increase in glycolysis in early hypoxia

To investigate mechanisms that sustain increased glycolysis in the absence of protein expression changes, we further explored the strong counter-correlation between aspartate and lactate levels. We attempted to prevent hypoxia-induced depletion of aspartate by providing cells with exogenous aspartate or its cell-permeable analogue, dimethyl-aspartate (DM-aspartate), to test whether the size of the intracellular aspartate pool influences lactate levels. Exogenous aspartate raised intracellular aspartate levels only modestly in normoxia, likely due to low expression of aspartate transporters ^71^ in MCF7 cells, and had minimal effects on intracellular aspartate and lactate concentrations when cells were exposed to 1% O_2_ for 3 h (**Figure S3A**). The reason DM-aspartate failed to have a more pronounced effect on intracellular aspartate levels is less clear. We therefore pursued a genetic strategy to test if preventing aspartate from decreasing in hypoxia interferes with the upregulation of glycolysis. CRISPR/Cas9-mediated knock-out of a known consumer of aspartate in the cytoplasm, GOT1, in MCF7 cells – henceforth referred to as GOT1ko cells – resulted in a 338 ± 45% increase in the steady-state levels of the intracellular aspartate pool (**Figure 3A-B**). Upon hypoxia treatment, the decrease in aspartate persisted in GOT1ko cells (**Figure 3B**), suggesting that GOT1 activity alone cannot account for the hypoxia-induced decrease in aspartate in wild-type cells. Further isotope labelling experiments indicated that the decrease in aspartate is likely due to attenuated production from glutamine but not from glucose (**Figure S3B-E**). Interestingly, the upregulation of glucose uptake and accumulation of both intracellular and secreted lactate after 3 h in hypoxia were markedly suppressed in GOT1ko cells (**Figure 3C-E**, **Figure S3F**). Glucose uptake remained attenuated also after 24 h in hypoxia in GOT1ko cells compared to wild-type cells (**Figure 3C**). Ectopic expression of HA-tagged GOT1 reversed the accumulation of aspartate in GOT1ko cells and restored hypoxia-induced lactate to levels similar to those in wild-type MCF7 cells (**Figure 3F-H**). These data suggested that GOT1 activity, rather than a decrease in aspartate concentration itself, is required to sustain the increase in glycolysis in early hypoxia.

**Figure 3.**
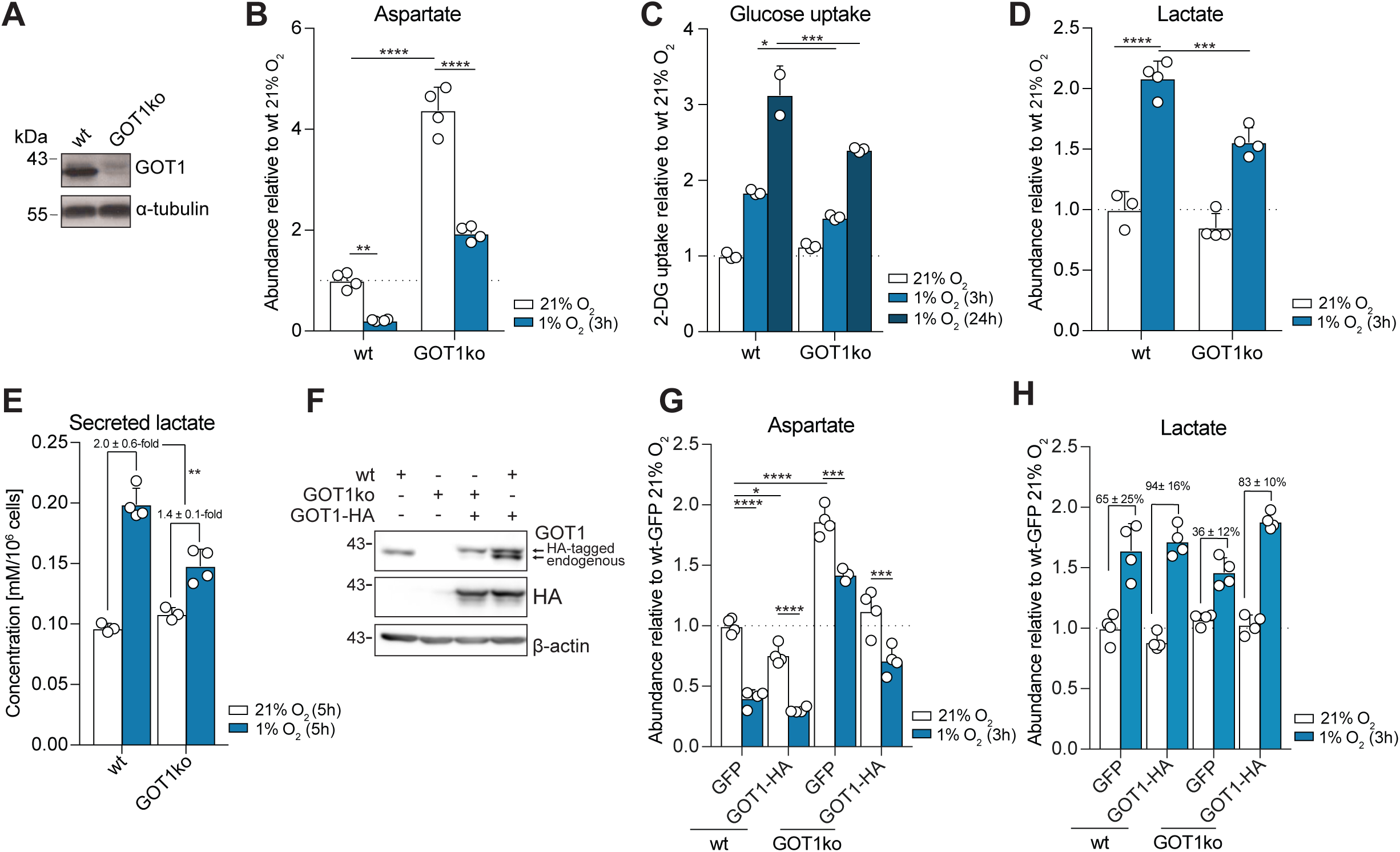
GOT1 supports increased glycolysis in early hypoxia. **A)** Western blot to assess levels of GOT1 in wild-type (wt) and GOT1ko MCF7 cells. **B)** Intracellular abundance of aspartate in wild-type (wt) and GOT1ko MCF7 cells incubated in 21% O_2_ or 1% O_2_ for 3 h. Data shown as mean ±SD (n = 4 cultures per condition) and significance was tested using two-way ANOVA with Tukey’s multiple comparisons test. **C)** Glucose (2DG) uptake of wild-type (wt) andand GOT1ko MCF7 cells in normoxia and after 3 and 24 h in 1% O_2_. Data shown as mean ±SD (n = 3 assays per condition) and significance was tested using two-way ANOVA with Sidak’s multiple comparison test. **D)** Intracellular abundance of lactate in wild-type (wt) andand GOT1ko MCF7 cells incubated in 21% O_2_ or 1% O_2_ for 3 h. Data shown as mean ±SD (n = 3-4 cultures per condition) and significance was tested using two-way ANOVA with Tukey’s multiple comparisons test. **E)** Lactate concentration in cell culture media of wild-type (wt) and GOT1ko MCF7 cells after 5 h incubation in 21% O_2_ or 1% O_2_. Data shown as mean ± SD (n = 3-4 cultures per condition). Statistical errors were propagated to calculate the error of the change in secreted lactate between normoxia and hypoxia for each cell line and significance was then tested using an unpaired t-test. Full time course shown in **Figure S3F**. **F)** Western blot to assess the levels of HIF1α, endogenous GOT1 and HA-tagged GOT1 in wild-type (wt) and GOT1ko MCF7 cells stably expressing GOT1-HA or GFP. **G-H)** Intracellular abundance of aspartate and lactate in wild-type (wt) and GOT1ko MCF7 cells stably expressing GOT1-HA or GFP at 21% O_2_ and after 3 h in 1% O_2_, relative to wild-type cells at 21% O_2_. Data shown as mean ± SD (n = 4 cultures per condition) and significance was tested using one-way ANOVA with Tukey’s multiple comparison test. For lactate, statistical errors were propagated to calculate the error of the change in lactate between normoxia and hypoxia for each condition and significance between these changes was then tested using an ordinary one-way ANOVA (not significant).

### GOT1 contributes to cytoplasmic NAD+/NADH balance by sustaining flux through MDH1

To probe the requirement of GOT1 for glycolysis, we quantified glycolytic and pentose phosphate pathway (PPP) intermediates by liquid chromatography-mass spectrometry (LC-MS). Metabolite pools upstream of GAPDH increased, while downstream metabolites decreased in GOT1ko compared to wild-type cells (**Figure 4A** and **Figure S4A**). This metabolic profile indicated a bottleneck for glycolytic flux at GAPDH ^5,72^, which was unlikely due to carbon substrate limitation since glucose uptake in normoxia was similar in both cell lines (**Figure 3C**). GAPDH activity depends on the availability of NAD^+^ and is attenuated by increased levels of NADH through competitive product inhibition ^73,74^. We therefore quantified NAD^+^ and NADH by LC-MS and compared the respective NAD^+^/NADH ratios in wild-type and GOT1ko cells. In wild-type cells, 3 h in 1% O_2_ led to a decrease in NAD^+^/NADH ratio from 6.0 ± 0.2 to 3.4 ± 0.2 (**Figure 4B**). In contrast, the NAD^+^/NADH ratio in GOT1ko cells was already lower (3.6 ± 0.4) at 21% O_2_ and did not change significantly upon incubation of cells in 1% O_2_. Notably, comparable NAD^+^/NADH ratios did not reflect equivalent redox states, because the hypoxia-induced decrease in NAD^+^/NADH ratio in wild-type cells was due to increased NADH levels, while the lower NAD^+^/NADH ratio in GOT1 cells reflected decreased NAD^+^ levels (**Figure 4B-D**). These findings are consistent with the idea that upregulation of glycolysis, associated with higher consumption of NAD^+^ by GAPDH, imposes an increased need for NADH oxidation to maintain redox balance. They also indicate that regeneration of NAD^+^ is compromised and may underlie impaired glycolysis in GOT1ko cells during early hypoxia.

**Figure 4.**
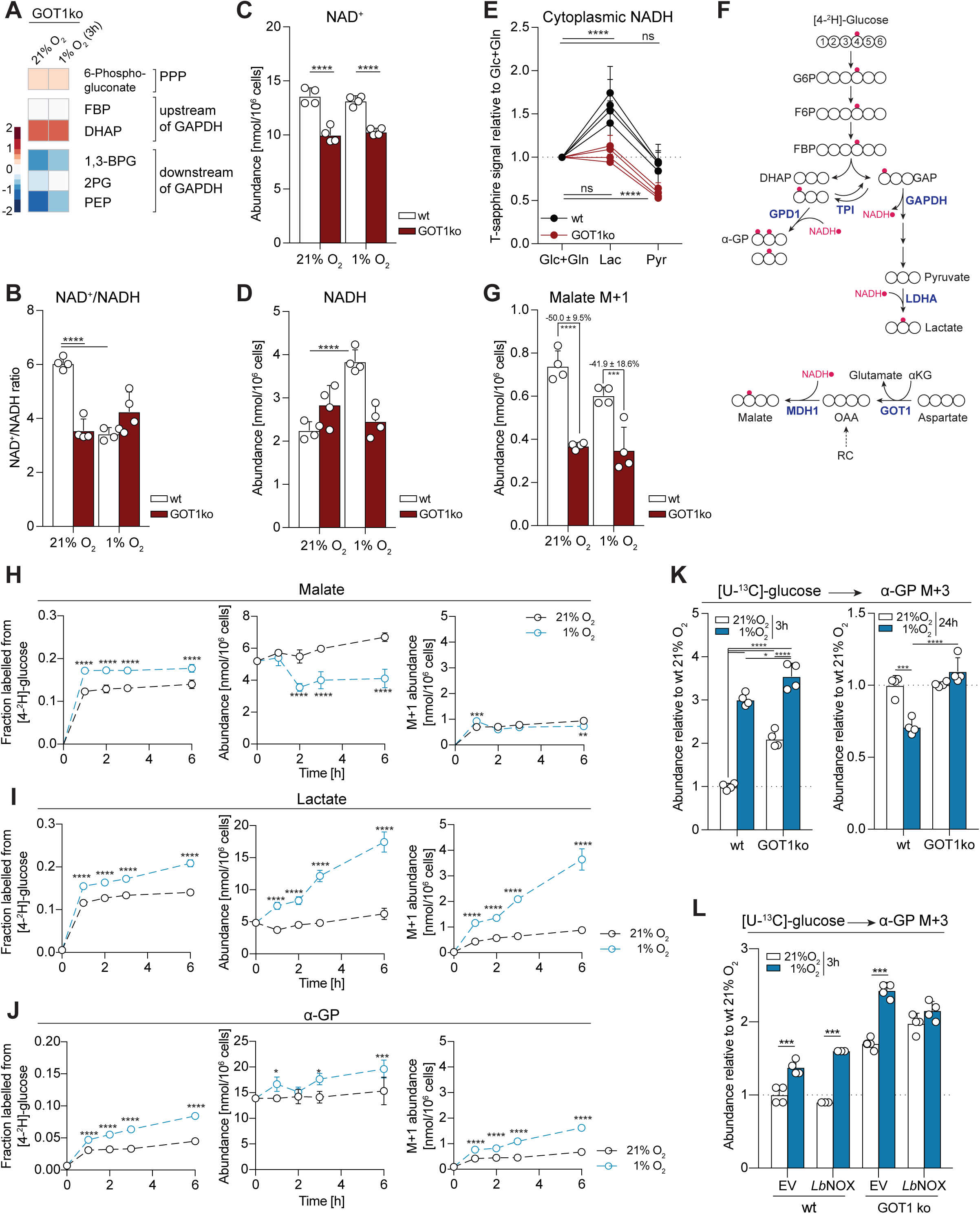
GOT1 supports MDH1 flux and cytoplasmic redox balance, but MDH1 flux does not change in hypoxia. **A)** Heatmap showing log_2_ fold-changes in the abundance of the indicated metabolites in GOT1ko at 21% O_2_ or after 3 h in 1% O_2_, compared to wild-type MCF7 cells under the same conditions (n = 4 cultures per time point and condition). Data for each condition separately are shown in **S4A**. **B-D)** NAD^+^/NADH ratio (**B**) calculated from the intracellular abundance of NAD^+^ (**C**) and NADH (**D**) in wild-type (wt) and GOT1ko MCF7 cells incubated in 21% O_2_ or 1% O_2_ for 3 h (n = 4 cultures per condition). Data are shown as mean ± SD and significance was tested using two-way ANOVA with Tukey’s multiple comparison test. **E)** Peredox T-sapphire fluorescence signal intensity of wild-type (wt) and GOT1ko MCF7 cells in buffer containing 5.5 mM glucose and 2 mM glutamine (Glc+Gln) and after sequential incubation first with 10 mM lactate (Lac) and then with 10 mM pyruvate (Pyr). Signal was normalized per nucleus and is shown relative to the Glc+Gln condition. Data points represent mean ± SD of 4 independent replicates per cell line (n = 25-55 cells per replicate) and significance relative to Glc+Gln was tested using one-way ANOVA with Dunnett’s multiple comparison test. See also **Figure S4B-C**. **F)** Schematic showing theoretical labelling patterns in the indicated metabolites from [4-^2^H]-glucose. Carbon atoms are shown in white and deuterium atoms are shown in red. Adapted from ^78^. **G)** Intracellular abundance of the M+1 isotopologues of malate in wild-type (wt) and GOT1ko MCF7 cells after incubation with [4-^2^H]-glucose for 3 h. Data shown as mean ± SD (n = 4 cultures per condition) and significance was tested using two-way ANOVA with Sidak’s multiple comparison test. See also **Figure S4E**. **H-J)** Fraction labelled from [4-^2^H]-glucose, absolute abundances and absolute abundances of M+1-labelled isotopologues from [4-^2^H]-glucose of the shown metabolites. Time points indicate duration of incubation at 21% O_2_ or 1% O_2_, as well as duration of incubation with the isotopic tracer. Data shown as mean ± SD (n = 4 cultures per condition) and significance was tested using two-way ANOVA with Sidak’s multiple comparison test. **K)** Intracellular abundance of α-glycerophosphate (α-GP) M+3 labelled from [U-^13^C]-glucose in wild-type (wt) and GOT1ko MCF7 cells after the indicated lengths of time in 21% O_2_ or 1% O_2_. Cells were incubated with the tracer for 3 or 24 h, respectively. Data shown as mean ±SD (n = 4 cultures per cell line and condition, experiment performed once) and significance was tested using two-way ANOVAs with Tukey’s multiple comparisons test. **L)** Intracellular abundance of α-glycerophosphate (α-GP) M+3 labelled from [U-^13^C]-glucose in wild-type (wt) and GOT1ko MCF7 cells, stably expressing an empty vector (EV) or *Lb*NOX. Cells were incubated with the tracer for 3 h in 21% O_2_ or 1% O_2_, respectively. Data shown as mean ± SD (n = 4 cultures per cell line and condition, experiment performed once) and significance was tested using two-way ANOVAs with Sidak’s multiple comparisons test.

The cellular pools of NAD(H) are asymmetrically distributed between subcellular compartments, due to the differential localisation of pyridine nucleotide precursors and biosynthetic pathways, as well as the impermeability of the inner mitochondrial membrane to NAD(H) ^75^. Since GAPDH activity depends on cytoplasmic NAD^+^/NADH, we specifically assessed cytoplasmic redox state in wild-type and GOT1ko cells using the genetically encoded NADH sensor Peredox ^76^. To this end, we recorded the basal Peredox T-sapphire signal of individual cells incubated in buffer supplemented with regular concentrations of the main carbon sources glucose and glutamine, and subsequently compared it to the T-sapphire signal after sequential incubation with only 10 mM lactate or 10 mM pyruvate. In the absence of extracellular glucose to counter-balance cytoplasmic redox changes, incubation of cells with lactate leads to production of NADH via LDHA and results in maximal Peredox T-sapphire signal that depends on the amount of available NAD^+^ ^76,77^. Conversely, incubation of cells with pyruvate leads to consumption of available cytoplasmic NADH via LDHA and minimises Peredox T-sapphire signal (**Figure S4B**). Assessment of the basal Peredox T-sapphire together with the maximal availability of NADH and NAD^+^ reported by incubation with pyruvate and lactate, respectively, allows comparison of the cytoplasmic redox state in wild-type versus GOT1ko cells.

In wild-type MCF7 cells, the basal Peredox signal was similar to the high NAD^+^/NADH state (Pyr), while lactate treatment (Lac) caused a dramatic increase in Peredox signal intensity irrespective of the order of substrate addition (**Figure 4E, Figure S4C**). Conversely, in GOT1ko cells extracellular lactate failed to increase cytoplasmic NADH, further supporting our interpretation of the cell population-level NAD^+^/NADH quantification by LC-MS that GOT1ko cells have a deficit in cytoplasmic NAD^+^. Accordingly, overexpression of GOT1-HA in wt cells enhanced the lactate-induced and suppressed the pyruvate-induced Peredox response; overexpression of GOT1-HA in GOT1ko cells restored the Peredox response to wt cell levels (**Figure S4D**). Together, these observations indicated that the attenuated increase in glycolysis of GOT1ko cells in early hypoxia may be due to decreased NAD^+^ availability.

GOT1 converts aspartate to oxaloacetate (OAA), a substrate of MDH1, which produces malate and concomitantly oxidises NADH to NAD^+^ (**Figure 4F**). To assess whether deletion of GOT1 influences MDH1 activity and thereby the production of cytoplasmic NAD^+^, we incubated cells with [4-^2^H]-glucose, which leads to production of cytoplasmic NAD^2^H that can be subsequently used by MDH1 to incorporate a deuterium into malate (malate M+1) ^78^ (**Figure 4F**). Malate M+1 abundance was 50 ± 10% and 42 ± 19% lower in normoxia and early hypoxia, respectively, in GOT1ko compared to wild-type cells (**Figure 4G**), while labelling of NADH was similar in both cell lines and conditions (**Figure S4E**). These results show that a significant fraction of MDH1 flux depends on GOT1 both in normoxia and in early hypoxia, consistent with the lower basal NAD^+^ levels we observed in GOT1ko cells (**Figure 4C**).

### MDH1 flux does not increase in early hypoxia

Other reports have previously indicated that increased flux through MDH1 is required to support glycolysis ^79^ and this may be associated with increased MDH1 expression ^80^. We also observed an increase in MDH1, but not GOT1 protein levels in hypoxia (**Fig S4F**), however, we found no significant differences in malate M+1 levels between normoxia and hypoxia in either cell line (**Figure 4G**). We therefore explored the contribution of GOT1, *versus* other potential OAA sources, to MDH1 flux in early hypoxia.

In wild-type MCF7 cells, the incorporation of deuterium into malate reached steady-state within 1 h in hypoxia (**Figure 4H**, left panel). After accounting for the significant decrease in the malate pool size (**Figure 4H**, middle panel), we found that the abundance of malate M+1 at steady-state was similar in 21% O_2_ and 1% O_2_ (**Figure 4H**, right panel). Malate M+1 levels remained relatively constant even after 6 h in hypoxia, despite the progressive decrease in aspartate over time (**Figure S4G**), indicating that aspartate did not become limiting for GOT1-MDH1 flux. Furthermore, overexpression of ectopic GOT1-HA in wild-type MCF7 cells modestly enhanced lactate accumulation (**Figure 3H**) suggesting that lower glycolysis under these conditions may be limited by GOT1 expression.

Reductive carboxylation (RC) has also been proposed as a source of OAA for MDH1 to support glycolysis in cells with mitochondrial defects or upon growth factor stimulation ^79,80^. Similar to previous reports ^81^, RC increased in MCF7 cells after 24 h in hypoxia, however, we found no significant increase in RC after 3 h in hypoxia (**Figure S4H**). GOT1ko cells showed a modest increase in RC (**Figure S4I**) but this was not sufficient to fully restore MDH1 flux (**Figure 4G**) and prevent the observed attenuation of glycolysis in early hypoxia (**Figure 3C**-**D**).

Together, these data showed that MDH1 flux is maximal in normoxia and does not further increase in early hypoxia.

### NAD^+^ is limiting for maximal flow of carbons from upper to lower glycolysis in early hypoxia

Given the lack of increase in MDH1 flux during early hypoxia, we investigated whether LDHA, which has an established role in supporting increased glycolysis in chronic hypoxia ^48,50^ has a similar role also in early hypoxia. In normoxia, the amount of lactate M+1 produced from [4-^2^H]-glucose was comparable to that of malate M+1 (**Figures 4H** and **4I**, right panels). In hypoxia, incorporation of ^2^H into lactate increased linearly over time, indicating an increased contribution of LDHA to NAD^+^ production (**Figure 4I**). We concluded that, in contrast to MDH1, flux through LDHA in normoxia is not maximal and can increase in early hypoxia despite the lack of increased LDHA protein expression.

Interestingly, we observed that production of α-glycerophosphate (α-GP, glycerol 3-phosphate) M+1 from [4-^2^H]-glucose also increased in hypoxia (**Figure 4J**), which was further reflected by the increased labelling of α-GP from [U-^13^C]-glucose (**Figure 4K**, wt cells on the left). α-GP is produced by α-GP dehydrogenase 1 (GPD1), from dihydroxyacetone phosphate (DHAP), which, alongside glyceraldehyde 3-phosphate (GAP) is a product of aldolase (**Figure S4J**). DHAP and GAP are interconverted by triose phosphate isomerase (TPI), which thermodynamically favours DHAP formation. However, in cells, high GAPDH activity rapidly consumes GAP and shifts the TPI equilibrium towards GAP formation, thereby allowing the reactions of lower glycolysis to occur ^73,82–86^. Therefore, increased incorporation of glucose carbons to α-GP may reflect a limitation in carbon flow from upper to lower glycolysis due to attenuated GAPDH activity, increased GPD1 activity, or both.

Consistent with this idea, GOT1ko cells, which have impaired NAD^+^-regenerating capacity (**Figure 4G**) and attenuated lower glycolysis (**Figure 4A**), show increased labelling of α-GP from [U-^13^C]-glucose compared to wild-type cells in normoxia (**Figure 4K**, left) despite lower glucose uptake (**Figure 3C**). Expression of the bacterial NADH oxidase *Lb*NOX ^87^ completely prevented the hypoxia-induced increase in α-GP labelling from [U-^13^C]-glucose in GOT1ko cells but not in wild-type cells (**Figure 4L**), further supporting our model that elevated efflux of glucose carbons to α-GP in early hypoxia when GOT1 is absent reflects limiting NAD^+^. Expression of exogenous GOT1-HA in GOT1ko cells decreased α-GP to similar levels as those found in parental cells in hypoxia and partly reversed the impairment in MDH1 flux (**Figure S4K**). Intriguingly, α-GP labelling decreased in wild-type cells after 24 h in hypoxia, compared to normoxia, but remained elevated in GOT1ko cells (**Figure 4K**, right). Together, these data suggested that, in early hypoxia, cellular NAD^+^-regenerating capacity is not sufficient for maximal flow of carbons from upper to lower glycolysis. After 24 h in hypoxia, decreased α-GP production indicates that flux through the reactions of lower glycolysis can match, or exceed, that of upper glycolysis, likely because of increased NAD^+^-regenerating capacity due to higher LDHA protein expression (**Figure 2A**) or increased RC via MDH1 (**Figure S4K**).

We next tested whether LDHA is sufficient for sustaining lower glycolysis or whether its function can be substituted by GOT1-MDH1. Knock-out of LDHA (LDHAko) in MCF7 cells effectively abrogated production of lactate from glucose and led to an accumulation of labelled pyruvate (**Figures 5A-B**), demonstrating that LDHA is the predominant LDH isoform in these cells. α-GP labelling from glucose increased in LDHAko cells, pointing to a bottleneck between upper and lower glycolysis. This interpretation was confirmed by the observed increase in the levels of metabolites in upper glycolysis and depletion of metabolites in lower glycolysis both under normoxia and, to a greater extent, under hypoxia (**Figure 5C** and **Figure S5A**). Notably, these changes in glycolytic intermediates were more pronounced than those in GOT1ko cells (**Figure 4A** and **Figure S4A**). This observation, combined with the enhanced synthesis of α-GP from glucose in LDHAko *versus* GOT1ko cells, revealed an increased reliance of lower glycolysis on LDH than GOT1-dependent MDH1 flux. Together, our data indicated that LDHA is necessary but not sufficient, even combined with GOT1-MDH1, to sustain maximal carbon flow from upper to lower glycolysis in hypoxia.

**Figure 5.**
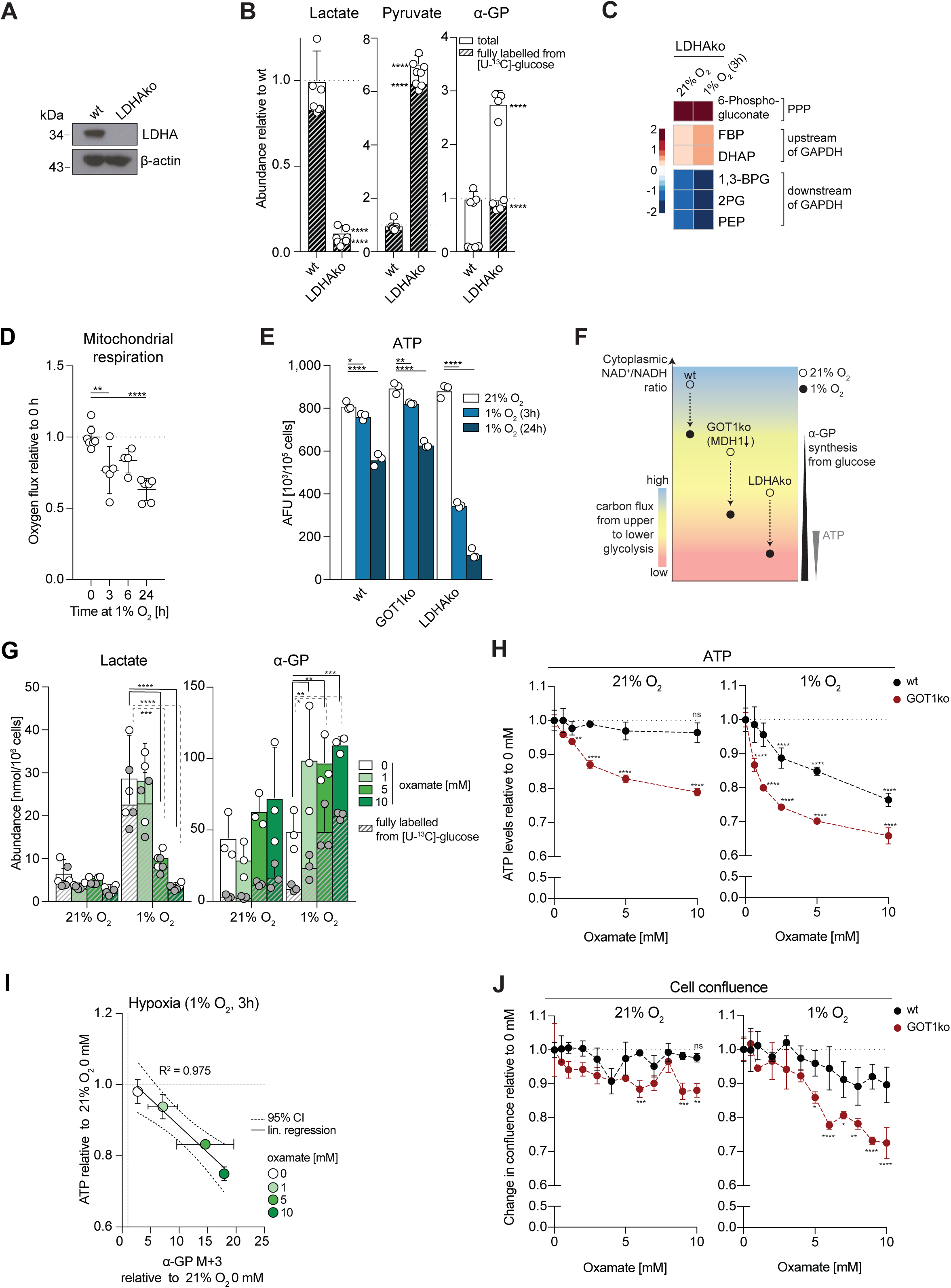
LDH has spare capacity in normoxia and is necessary to maintain ATP levels in early hypoxia. **A)** Western blot to assess the levels of LDHA in wild-type (wt) and LDHAko MCF7 cells. **B)** Intracellular abundance of pyruvate, lactate and α-glycerophosphate (α-GP) in wild-type (wt) and LDHAko MCF7 cells. Striped bars represent the fraction of metabolites fully labelled from [U-^13^C]-glucose after 3 h incubation with the tracer. Data shown as mean ± SD (n = 4 cultures per condition) and significance compared to wild-type cells was tested using two-way ANOVA with Sidak’s multiple comparison test. **C)** Heatmap showing log_2_ fold-changes in the abundance of the indicated metabolites in LDHAko at 21% O_2_ or after 3 h in 1% O_2_, compared to wild-type MCF7 cells in the same conditions (n = 4 cultures per time point and condition). Data for each condition separately are shown in **S5A**. Wild-type data are the same as shown in **Figure 4A and S4A** and statistical tests were performed on the whole data set. D) Mitochondrial respiration of MCF7 cells after incubation at 1% O_2_ for the indicated lengths of time. Cellular oxygen consumption was corrected for ROX (residual oxygen consumption) by addition of the complex III inhibitor antimycin A. Data shown as mean ± SD (n = 4-7 replicates per condition; graphs show combined replicates of two independent experiments) and significance compared to 0 h was tested using one-way ANOVA with Dunnett’s multiple comparison test. **E)** ATP levels in wild-type (wt), GOT1ko and LDHAko MCF7 cells at 21% O_2_ and after 3 h or 24 h in 1% O_2_. Data shown as mean ± SD (n = 3 assays per cell line condition) and significance relative to 21% O_2_ was tested ANOVA with Dunnett’s multiple comparison test. See also Figure S5B. **F)** Schematic summarizing the observed effects of GOT1 and LDHA deletion on carbon flux from upper to lower glycolysis (indicated by the color scale) and changes in cellular ATP during early hypoxia. Decreased NAD^+^/NADH ratio in GOT1ko cells leads to an attenuation of carbon flux into lower glycolysis that is not large enough to affect ATP levels. In contrast, loss of LDHA leads to more profound inhibition of lower glycolysis, associated with ATP depletion and cell death. wt: wild-type cells. **G)** Intracellular abundance of lactate and α-glycerophosphate (α-GP) in MCF7 cells treated with a range of concentrations of the LDHA inhibitor oxamate for 3 h at 21% O_2_ or 1% O_2_. Striped bars represent the fraction of metabolites fully labelled from [U-^13^C]-glucose after 3 h incubation with the tracer. Data shown as mean ± SD (n = 3 cultures per condition) and significance was tested using two-way ANOVA with Dunnett’s multiple comparison test. **H)** ATP levels in wild-type (wt) and GOT1ko MCF7 cells at 21% O_2_ and after 3 h in 1% O_2_ treated with the indicated oxamate concentrations for 3 h. Data shown as mean ± SD (n = 3 assays per cell line and condition) and significance relative to 0 mM was tested ANOVA with Dunnett’s multiple comparison test. **I)** Scatter plot showing changes in α-glycerophosphate (α-GP) M+3 labelled from [U-^13^C]-glucose (3 h incubation with the tracer) relative to control cells in normoxia versus the changes in ATP levels relative to control cells in normoxia in MCF7 cells treated with a range of oxamate concentrations for 3 h at 1% O_2_. **J)** Change in cell confluence of wild-type (wt) and GOT1ko MCF7 cells within 24 hours at 21% O_2_ or 1% O_2_ with the indicated oxamate concentration in cell culture media, shown relative to 0 mM oxamate per cell line. Data shown as mean ± SD (n = 3 cultures per cell line and condition) and significance relative to wt cells was tested using two-way ANOVA with Sidak’s multiple comparison test.

### GOT1 and LDHA synergistically maintain ATP homeostasis and cell survival in hypoxia

An important function of increased glycolysis in chronic hypoxia, when respiration is suppressed, is to maintain intracellular ATP levels ^8^. We observed a comparable decrease in respiration of wt cells at 3 and 24 h hypoxia (23 ± 18 % vs 37 ± 11 %, respectively, **Figure 5D**), that was accompanied by a more pronounced decrease in ATP after 24 h (31 ± 4%) than after 3 h in hypoxia (6 ± 2%, **Figure 5E**). These results suggested that increased glycolysis could also preserve ATP levels upon suppression of mitochondrial respiration in early hypoxia.

As our data pointed to a differential role for GOT1-MDH1 and LDHA in sustaining lower glycolysis, we compared the relative ability of GOT1ko and LDHAko cells to maintain ATP homeostasis. ATP levels in both GOT1ko and LDHAko cells were comparable to those in wild-type cells in normoxia (**Figure 5E**), suggesting that remaining flux through lower glycolysis after deletion of either GOT1 or LDHA is sufficient to maintain ATP homeostasis in normoxia. Under hypoxia, ATP levels decreased similarly in wild-type and GOT1ko cells, but more significantly in LDHAko cells, both at 3 h and 24 h (**Figure 5E**). Expression of low levels of exogenous LDHA attenuated the hypoxia-induced decrease in ATP from 56% to 32% (**Figure S5B**). ATP depletion in LDHAko cells was accompanied by a 68 ± 8% loss of cell mass after 24 h in hypoxia compared to normoxia, whereas, under the same conditions, wild-type cells showed a more modest decrease in cell mass (15 ± 4%, **Figure S5C**). These data are in line with the increased reliance of lower glycolysis on LDHA compared to GOT1-MDH1 in early hypoxia and indicate that MDH1 fuelled by OAA from GOT1, or other sources, cannot compensate for decreased LDHA activity. Conversely, our results suggest that LDHA may suffice to support ATP production upon loss of GOT1 (**Figure 5F**).

To test the dependence of GOT1ko cells on LDHA for maintaining ATP levels, we treated GOT1ko cells with oxamate, a competitive inhibitor of LDHA ^88^ that led to a dose-dependent decrease in the production of lactate from [U-^13^C]-glucose (**Figure 5G**, left). In agreement with our findings in LDHAko cells, treatment with oxamate had no effect on ATP levels in wild-type cells in normoxia but led to a dose-dependent decrease in ATP levels in hypoxia (**Figure 5H**), without affecting mitochondrial respiration (**Figure S5D**). The oxamate-induced decrease in ATP under hypoxia correlated remarkably well (R^2^ = 0.9754) with the magnitude of the increase in α-GP synthesis from glucose under the same conditions (**Figure 5G**, right and **5I**), further supporting our model that increased α-GP synthesis reflects efflux of glucose carbons from glycolysis via GPD1 and diminished activity of lower glycolysis. In contrast to untreated GOT1ko cells, treatment of GOT1ko cells with oxamate resulted in lower ATP levels even in normoxia. This decrease in ATP was further exacerbated after 3 h in hypoxia (**Figure 5H**) but did not occur either in normoxia or hypoxia with oligomycin (**Figure S5E**). Furthermore, oxygen consumption in GOT1ko cells was modestly decreased compared to that of wild-type cells (**Figure S5F**). Together, these observations indicated that GOT1ko cells rely more on glycolytic flux supported by LDHA, rather than compensatory mitochondrial respiration, to maintain intracellular ATP levels. Importantly, oxamate caused a more profound decrease in GOT1ko than wild-type cell number in hypoxia than in normoxia and this decrease was rescued by exogenous GOT1-HA (**Figure 5J** and **S5G**). Collectively, these data revealed that while GOT1-MDH1 and LDHA contribute differentially to ATP production from lower glycolysis, they synergise to maintain ATP homeostasis in early hypoxia and contribute to cell survival after 24 h in hypoxia.

In summary, our results suggest that enhanced upper glycolysis in early hypoxia imposes an increased need for NAD^+^ to sustain maximal flow of carbons to lower glycolysis. This increased requirement for NAD^+^ is met by reserve LDHA capacity and is further supported by (saturated) GOT1-MDH1, to sustain ATP homeostasis in early hypoxia. However, even with combined action of LDHA and MDH1, NAD^+^ regeneration is not enough to achieve maximal flow of carbons from upper to lower glycolysis.

### GOT1 consumes αKG to attenuate PHD activity and promote HIF1α stabilisation

To investigate whether loss of GOT1 also affects the long-term hypoxic response, we first monitored wild-type and GOT1ko cell proliferation over 2 days. We found that loss of GOT1 did not affect proliferation either in normoxia or hypoxia (**Figure S6A**). We next compared gene expression in wild-type and GOT1ko cells incubated in 1% O_2_ for 24 h. Both cell lines exhibited wide-spread gene expression changes in hypoxia, compared to normoxia, including changes in HIF1α target genes (**Figure S6B**). However, the induction of HIF1α target gene expression was markedly suppressed in GOT1ko cells compared to wild-type cells (**Figure 6A**). This suppression was not due to a defect in transcription, as the profile of global gene expression changes induced by hypoxia in GOT1ko cells was largely similar to that of wild-type cells (**Figure S6C**). Therefore, these data show that HIF1α-dependent transcription is attenuated in GOT1ko cells.

**Figure 6.**
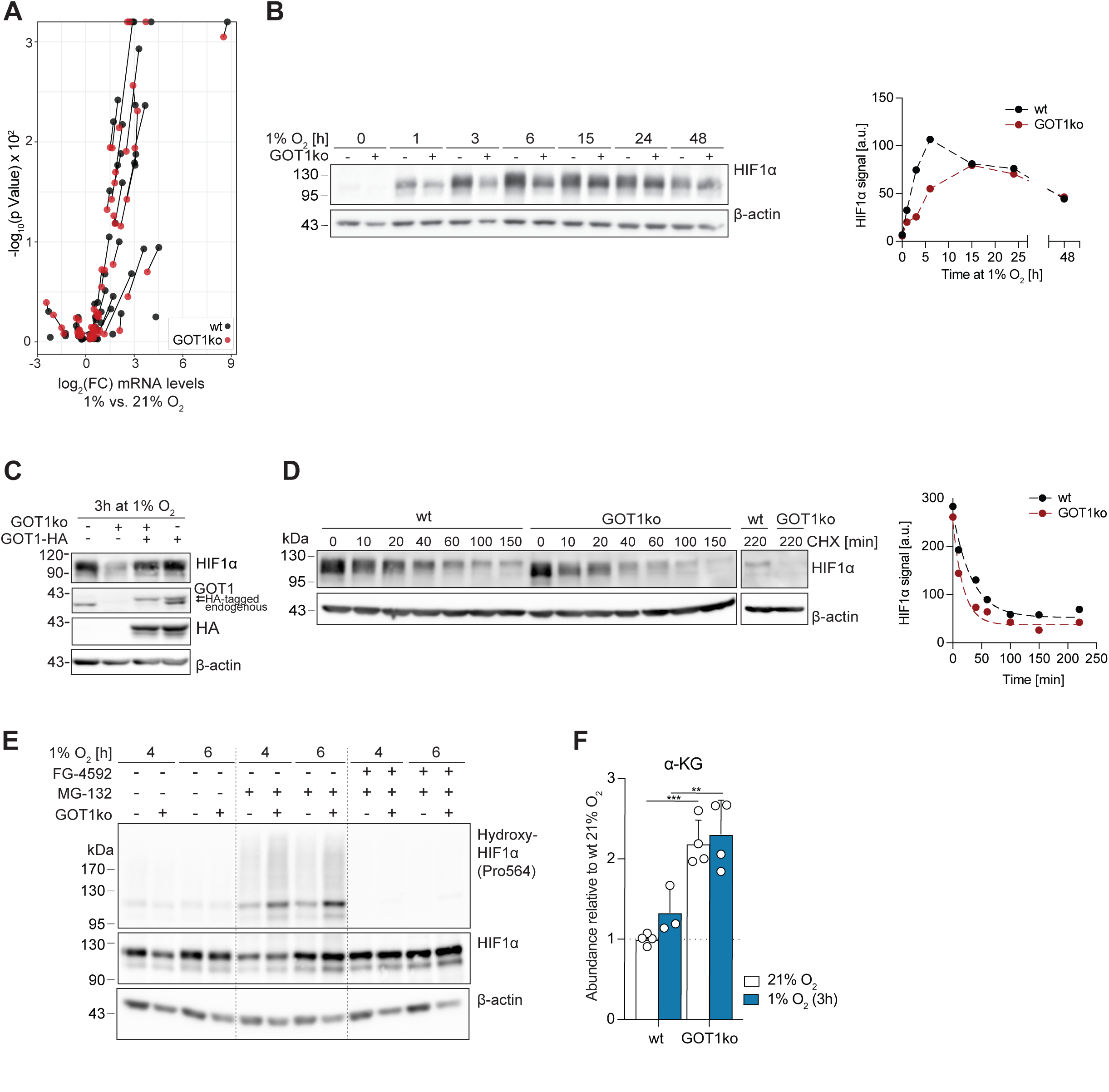
Elevated αKG levels, increased HIF1α hydroxylation and attenuated HIF1α stabilisation in GOT1ko cells under hypoxia. **A)** Volcano plot of gene expression changes of a panel of HIF1α target genes in wild-type (wt) MCF7 and GOT1ko cells exposed to 1% O_2_ for 24 h, compared to control cells in normoxia (n = 3 cultures per condition; only changes with FDR<0.01 are shown). Lines connect identical genes in the two cell lines to illustrate the differences in hypoxia-induced gene expression changes. **B)** Western blot to assess HIF1α protein levels in wild-type (wt) and GOT1ko MCF7 cells exposed to 1% O_2_ for the indicated lengths of time. The graph on the right shows the quantification of the HIF1α signal. Graph shows one representative experiment, performed three times. **C)** Western blot to assess the protein levels of HIF1α, endogenous GOT1 and HA-tagged GOT1 in wild-type (wt) and GOT1ko MCF7 cells stably expressing GOT1-HA or GFP and exposed to 1% O_2_ for 3 h. **D)** Western blot to assess HIF1α protein levels in wild-type (wt) and GOT1ko MCF7 cells exposed to 1% O_2_ for 3 h and then treated with cycloheximide (CHX, 20 μM) for the indicated lengths of time. Graph on the right shows the quantification of the HIF1α signal. Graph shows one representative experiment, performed twice. **E)** Western blot to assess the levels of HIF1α, and HIF1α hydroxylated at proline 564 (Pro564) in wild-type (wt) and GOT1ko MCF7 cells exposed to 1% O_2_ for the indicated lengths of time. Cells were treated with the PHD inhibitor FG-4592 (50 μM), the proteasome inhibitor MG-132 (10 μM) or a combination of both for the duration of the experiment. See also **Figure S6F** for additional controls. **F)** Intracellular abundance of α-ketoglutarate (α-KG) in wild-type (wt) and GOT1ko MCF7 cells after 3 h at 21% O_2_ or 1% O_2_. Data shown as mean ±SD (n = 4 cultures per condition) and significance was tested using two-way ANOVA with Sidak’s multiple comparison test.

Decreased induction of HIF1α target mRNAs was associated with both a delay and decrease in the hypoxia-induced accumulation of HIF1α protein in GOT1ko cells (**Figure 6B**). This was particularly evident within the first hours in hypoxia, whereas the kinetics of the decrease in HIF1α protein levels at longer times (>15 h) under hypoxia ^89^ were similar. Re-expression of HA-tagged GOT1 restored HIF1α expression, showing a GOT1-specific effect (**Figure 6C**). These data indicate that GOT1 promotes HIF1α stabilisation in early hypoxia and suggest that attenuation of HIF1α target gene expression in GOT1ko cells in later hypoxia reflects a cumulative effect of suppressed early HIF1α stabilisation.

Although we observed a modest decrease (<21%) in HIF1α mRNA levels in GOT1ko cells compared to wild-type cells (**Figure S6D**), treatment of cells with the proteasome inhibitor MG-132 led to similar kinetics of HIF1α protein accumulation in both cell lines (**Figure S6E**), indicating that decreased protein synthesis was unlikely to be the cause for the difference in HIF1α protein levels. We therefore reasoned that the difference in HIF1α stabilisation between wild-type and GOT1ko could be attributable to higher rates of HIF1α degradation in GOT1ko cells. When, after 3 h in hypoxia, HIF1α protein translation was inhibited with cycloheximide, HIF1α protein levels decreased more rapidly in GOT1ko cells than in wild-type MCF7 cells (**Figure 6D**). Furthermore, treatment with MG-132 eliminated the difference in HIF1α levels between wild-type and GOT1ko cells but revealed higher levels of hydroxylated HIF1α in GOT1ko than in wild-type MCF7 cells (**Figure 6E, Figure S6F**). The HIF1α hydroxylation signal was eliminated after treatment with the PHD inhibitor FG-4592, confirming that increased HIF1α hydroxylation is due to PHD activity. Taken together, these data suggest that GOT1ko cells retain higher PHD activity in hypoxia that could account for the delay in HIF1α stabilisation.

Although we observed small differences in mRNA expression of PHDs in GOT1ko compared to wild-type MCF7 cells (**Figure S6D**), such differences were not reflected on the protein level (**Figure S6G**) and pointed to increased PHD activity. PHD activity depends on O_2_ which, binds to PHDs in an αKG-dependent manner, therefore fluctuations in both O_2_ and αKG can influence PHD activity. αKG abundance increased in GOT1ko cells compared to wild-type cells (**Figure 6F**), consistent with αKG being a substrate of GOT1. These data indicate that, in addition to its immediate contribution to glycolysis upon oxygen limitation, GOT1 activity contributes to αKG turnover and thereby controls the kinetics of HIF1α stabilisation.

## DISCUSSION

Adaptation to low oxygen is critical for the survival and proliferation of cancer cells ^90^. While the PHD-HIF1α signalling axis is a key orchestrator of cellular responses to chronic hypoxia, the mechanisms that allow cells to survive until a full HIF1α response is established are not well understood. In this study, we present evidence that GOT1 primes the response of cells to oxygen limitation in two ways: firstly, it provides OAA used by MDH1 to produce NAD^+^ that supports increased glycolysis in early hypoxia; and, secondly, it maintains low intracellular αKG levels, which, in combination with low oxygen, attenuate PHD activity and thereby lead to efficient HIF1α stabilisation. Functionally, through this dual role, GOT1 supports glycolytic ATP production in the short term, as well as robust long-term HIF1α target gene expression (**Figure 7**).

**Figure 7.**
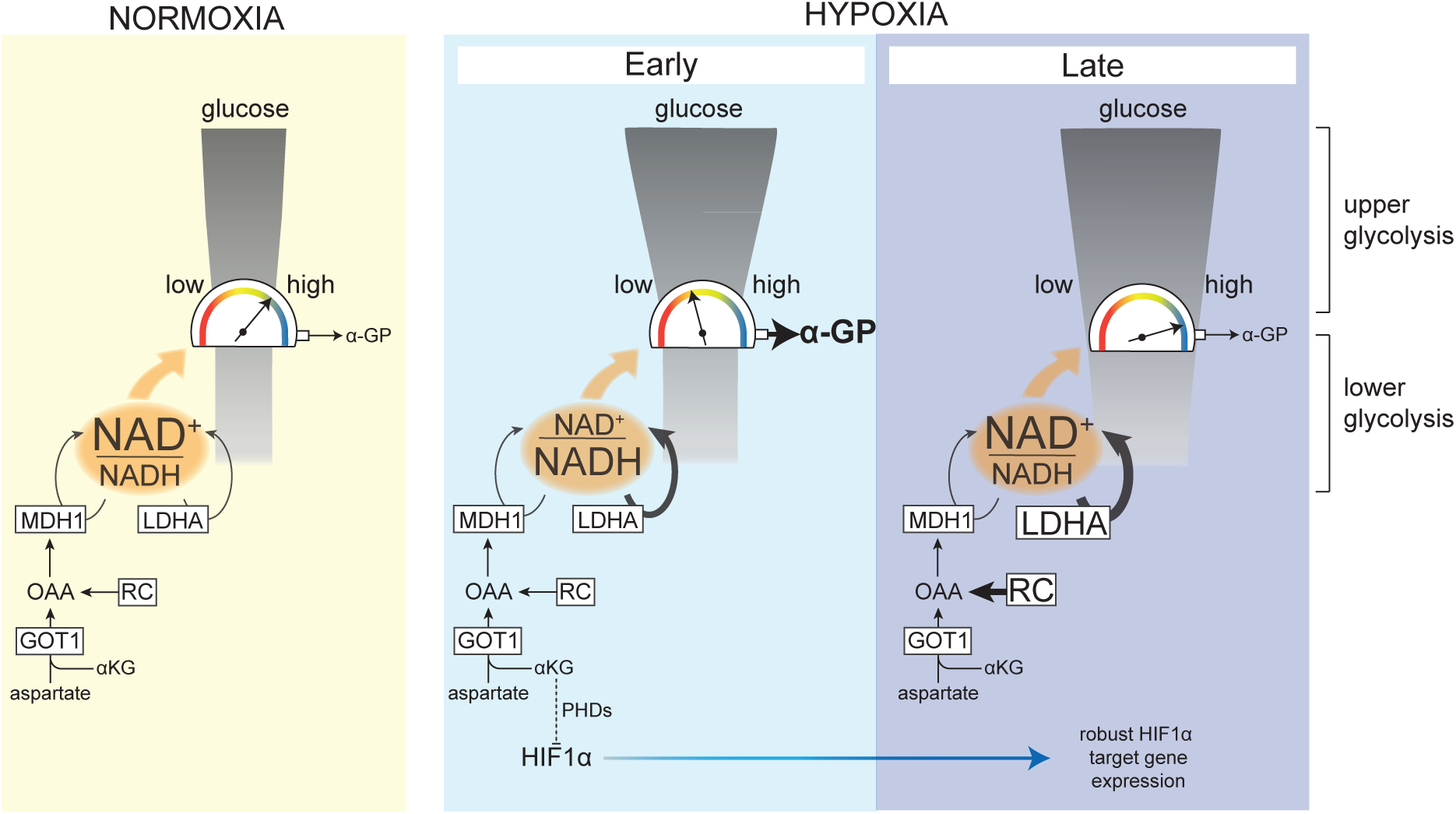
Model summarizing the dual role of GOT1 in enabling the cellular response to hypoxia. In **normoxia**, carbon flux through lower glycolysis matches that of upper glycolysis because LDHA and GOT1-driven MDH1 provide sufficient NAD^+^, which is needed for the flow of carbons (indicated by the *high* reading of the gauge) to lower glycolysis. The colored scale for the reading of the gauge indicates flux from upper to lower glycolysis. In **early hypoxia**, elevation of upper glycolysis increases the requirement for regeneration of NAD^+^, which is supported by an increase in the flux through LDHA and by GOT1-dependent MDH1 activity that does not increase compared to normoxia. However, carbon flow to lower glycolysis is limited by NAD^+^ in early hypoxia, as indicated by the increased efflux of glucose carbons to α-GP. In **late hypoxia**, increased RC provides additional OAA for MDH1 and, combined with increased LDHA expression, confers additional NAD^+^-regenerating capacity enabling increased flow of carbons to lower glycolysis. In parallel, GOT1 consumes αKG (an essential co-factor for PHDs), which, in combination with lower oxygen, suppresses HIF1α hydroxylation and therefore promotes its stabilization, leading to robust HIF1α target gene expression later in hypoxia. RC: reductive carboxylation.

It has long been recognised that oxygen suppresses glycolysis in both healthy and transformed cells, a phenomenon known as the Pasteur effect. Accordingly, we find that, upon oxygen limitation, glycolysis increases within 3 h and demonstrate that this occurs in a HIF1α-independent manner. The strong counter-correlation between aspartate and lactate levels led us to investigate how aspartate metabolism is linked to glucose metabolism. These investigations revealed that knock-out of GOT1 resulted in decreased flux through MDH1 and caused a lower NAD^+^/NADH ratio. Furthermore, loss of GOT1 attenuated lactate production from glucose only in hypoxia, pointing to an increased need for MDH1-derived NAD^+^ selectively under oxygen-limiting conditions.

GOT1 comprises one of the enzymes in the MAS (malate-aspartate shuttle), a system of proteins that translocates electrons between cytosolic and mitochondrial NADH. In chronic hypoxia, MAS activity has been broadly thought to be attenuated as a consequence of lower mitochondrial respiration or decreased production of aspartate, the substrate of GOT1 ^18,91–93^, in a HIF1α-dependent manner ^94^. In early hypoxia, we showed that both mitochondrial respiration (as previously shown ^95^) and aspartate synthesis from glutamine decreased. Therefore, our finding that the aspartate-consuming GOT1 is required for increased glycolysis in early hypoxia was, at first sight, paradoxical. However, we show that the amount of labelled malate produced from [4-^2^H]-glucose was similar in normoxia and hypoxia. This finding suggests that MDH1 flux is not impaired, even when aspartate levels decrease to less than 30% of those in normoxic cells, unlike the limitation in biomass production due to low aspartate seen in cells in chronic hypoxia or with mitochondrial defects ^71,91,96^. Importantly, overexpression of GOT1 led to increased lactate levels in hypoxia (**Figure 3H**). Together, these results support the idea that GOT1 expression, rather than aspartate levels, is limiting for MDH1-produced NAD^+^ and indicate that MDH1 flux in normoxia (as in other proliferating cells ^97^) and early hypoxia is, effectively, saturated.

Various well-established allosteric and signalling mechanisms that increase flux through the first enzymatic steps in glycolysis and glucose transport lead to increased upper glycolysis in hypoxia ^55,56,58,63–65,98,99^. It is, therefore, reasonable to suggest that such mechanisms elevate the activity of upper glycolysis in early hypoxia and impose an increased requirement for NAD^+^ that is used by GAPDH to enable the flow of incoming carbons to lower glycolysis. Under this light, the lack of increased MDH1 flux prompted us to explore other mechanisms that may regenerate NAD^+^ to support elevated glycolysis in early hypoxia.

In contrast to MDH1, lactate labelling from [4-^2^H]-glucose increased in early hypoxia. In the absence of detectable changes in protein levels, this result suggested that LDHA has reserve capacity in normoxia that can be used in early hypoxia to sustain NAD^+^. Intriguingly, we also observed increased α-GP synthesis from glucose in early hypoxia, which indicated a shift of the TPI equilibrium towards DHAP, the substrate of the α-GP–producing enzyme GPD1. The TPI equilibrium is influenced by the relative activities of GAPDH (which depends on NAD^+^ – see scheme in **Figure S4J**) and GPD1 (which depends on NADH). In hypoxia, a lower NAD^+^/NADH ratio, reflecting increased NADH levels, could either inhibit GAPDH ^73^, promote GPD1 activity ^100^ or both. α-GP synthesis increased following the knock-out of GOT1 or LDHA, both conditions that led to decreased lower glycolysis. Importantly, the LDHA inhibitor oxamate did not decrease respiration, confirming that the observed changes in α-GP are due to increased production from glucose, rather than decreased consumption due to lower activity of GPD2, an enzyme that converts α-GP to DHAP and provides reduced flavin adenine dinucleotide (FADH_2_) for mitochondrial respiration. Furthermore, increased α-GP synthesis after treatment of cells with oxamate correlated well with the decrease in ATP levels under the same conditions, which is likely due to attenuated lower glycolysis given that mitochondrial respiration was unchanged. Together, these observations are in line with a model where efflux of glucose carbons from the core glycolytic pathway to α-GP reflects a bottleneck at the GAPDH step.

Consequently, increased α-GP in early hypoxia strongly suggests that NAD^+^ provided by LDHA, even when supplemented by basal GOT1-MDH1 activity and possibly other pathways that support glycolysis in chronic hypoxia ^101^, is not sufficient for the increased amount of glucose carbons from upper glycolysis to flow into lower glycolysis (**Figure 7**–early hypoxia). Importantly, a model where increased upper glycolysis due to the Pasteur effect overwhelms GAPDH capacity also elucidates the *apparent* increase in the reliance of glycolysis on GOT1-MDH1 in hypoxia, even though flux through this pathway is not elevated (**Figure 7**). In normoxia, a lower amount of incoming carbons from upper glycolysis can be sustained by sub-saturated LDHA and saturated GOT1-MDH1. However, upon elevation of upper glycolysis in early hypoxia, a greater need for NAD^+^ arises, requiring “all hands on deck” to provide as much NAD^+^ as possible, therefore increasing the apparent reliance on GOT1-MDH1. As a consequence, combined inhibition of LDHA and GOT1 is detrimental to cells only in hypoxia but not in normoxia (**Figure 5J**), an effect that is associated with an ATP deficit (**Figure 5H**). It is likely that, for long-term survival in hypoxia, GOT1-supported cellular bioenergetics work together with other functions of MAS components shown to sustain biomass production in cells subjected to chronic hypoxia or with mitochondrial deficits ^71,91,96^.

In addition to GOT1, another major source of cytoplasmic OAA for MDH1 is ATP citrate lyase (ACL). In chronic hypoxia, RC of glutamine can provide carbons for lipids and OAA via ACL. RC-derived OAA supports MDH1-dependent NAD^+^ generation, which is required for glycolysis in cells stimulated with growth factors for 24 h ^80^ and in cells that harbour stable genetic mutations causing mitochondrial dysfunction ^79^. In the latter case, increased RC is caused by a decreased NAD^+^/NADH ratio. We also found a significant decrease in the NAD^+^/NADH ratio after 3h under hypoxia, however RC during this time remained unchanged. Nevertheless, RC increased after 24 h, consistent with previous reports that RC is controlled by HIF1α ^81,102^. Furthermore, a modest increase in RC in GOT1ko cells did not suffice to rescue the attenuation of glycolysis upon loss of GOT1.

Interestingly, cells with profound mitochondrial defects take up more aspartate from the media compared to isogenic cells with more functional mitochondria, and exogenous aspartate increases lactate in a GOT1-dependent manner ^79^. However, whether metabolism of endogenous aspartate via GOT1 itself is sufficient for glycolysis, and the relative contribution of GOT1 and RC to glycolysis remained unexplored. Intriguingly, higher degree of mitochondrial dysfunction is associated with decreased flux through the MAS suggesting that RC, rather than GOT1, may have a more important role for sustaining glycolysis in cells harbouring mitochondrial defects.

Our observation that, in early hypoxia, glycolytic flux is not maximal, even when supported by both LDH and GOT1-MDH1, elucidates the need for increased LDHA expression and increased RC to sustain MDH1 in chronic hypoxia. Such a model is further corroborated by our observation that α-GP levels decrease after 24 h in hypoxia, when both LDH expression and RC are increased, and are therefore expected to bestow a higher cellular capacity to regenerate glycolytic NADH (**Figure 7**–late hypoxia). Combined with evidence from the studies discussed above, our findings highlight the possibility that the mechanisms employed to provide NAD^+^ to glycolysis, and their relative contributions, may vary depending on different cues (hypoxia, growth factor stimulation, mitochondrial dysfunction) and the different lengths of time they require to increase glycolysis. It is conceivable that combinatorial targeting of multiple redox pathways, as we and others ^80^ showed, may be a useful therapeutic strategy to attenuate glycolysis. However, successful selection of relevant target combinations should rely on the relative contributions of these systems to sustaining glycolysis depending on the cellular and physiological context.

Our investigations into the long-term effects of GOT1 knock-out also pointed to a role for GOT1 in influencing HIF1α protein levels. While many αKG-consuming and -producing enzymes likely contribute to determining the steady-state intracellular concentration of αKG ^103,104^, our results suggest that GOT1 is a significant consumer of αKG. Given the critical role for αKG in PHD activity, increased αKG in GOT1ko cells is associated with decreased protein stability. Although HIF1α expression is not completely suppressed, a delay in stabilisation suffices to attenuate HIF1α target gene transcription in the long term, likely due to a cumulative effect over a period of several hours.

Chronic hypoxia develops over periods of tumour growth that are longer than the lengths of hypoxia treatment we used in our experiments. Nevertheless, hypoxia-reoxygenation cycles (also referred to as intermittent hypoxia) have important functional consequences for both tumour physiology and response to therapy ^105–113^ and occur in timescales that range between minutes and hours ^114–116^. Therefore, elucidating what factors determine the ability of cells to rapidly adapt during intermittent hypoxia is crucial. There is significant evidence that other cellular responses to low oxygen, such as suppression of translation ^117^ and increased ROS production ^118^, can occur within minutes to hours and also play an important role for cell survival. Our findings suggest that GOT1 functions help maintain cells in a primed state that increases their chances of survival when oxygen becomes limiting and, more broadly, support the notion that, upon exposure to stress, cells employ specific mechanisms that help them survive while other adaptive processes, that require more time, are established.

## ACKNOWLEDGMENTS

We would like to thank all members of the Anastasiou lab for valuable discussions and input throughout this work. We thank Mike Howell (Crick High Throughput Screening Science Technology Platform) for advice and help with cell proliferation and viability measurements; Hiroshi Kondo, Erik Sahai (Crick Tumour Cell Biology Laboratory) and Kurt Anderson (Crick Advanced Light Microscopy STP) for help and advice with microscopy-based NADH measurements; the staff at the Crick Advanced Sequencing STP for help with RNA sequencing. We acknowledge Sean O’Callaghan (Bio21 institute, The University of Melbourne) for the algorithm to correct for natural isotope abundance in metabolomics data. We also thank Salihanur Darici for technical help, and Alex Gould and Pia Ballschmieter for critical reading of the manuscript. pMSCV-Peredox-mCherry-NLS was a gift from Gary Yellen (Addgene plasmid # 32385), SpCas9(BB)-2A-Puro (PX459) V2.0 was a gift from Feng Zhang (Addgene plasmid # 62988). pUC57-*Lb*NOX was a gift from Vamsi Mootha (Addgene plasmid # 75285). Funding for this work was provided to D.A. by the MRC (MC_UP_1202/1) and by the Francis Crick Institute which receives its core funding from Cancer Research UK (CC2113), the UK Medical Research Council (CC2113) and the Wellcome Trust (CC2113). For the purpose of Open Access, the authors have applied a CC-BY public copyright licence to any Author Accepted Manuscript version arising from this submission.

## AUTHOR CONTRIBUTIONS

FG designed and performed experiments, analysed and interpreted data, and contributed to the writing of the manuscript; AA contributed to metabolomics and redox biosensor experiments; AJ and PN performed cellular respiration experiments; MSdS and JIM assisted with, and advised on, metabolomics experiments and method development; JK performed RNA-seq data analysis; SG helped with microscopy; LF generated constructs and helped with the characterisation of cell lines; DA conceived and supervised the study, designed experiments, interpreted data and wrote the manuscript. All authors reviewed and commented on the manuscript.

## Key resources table

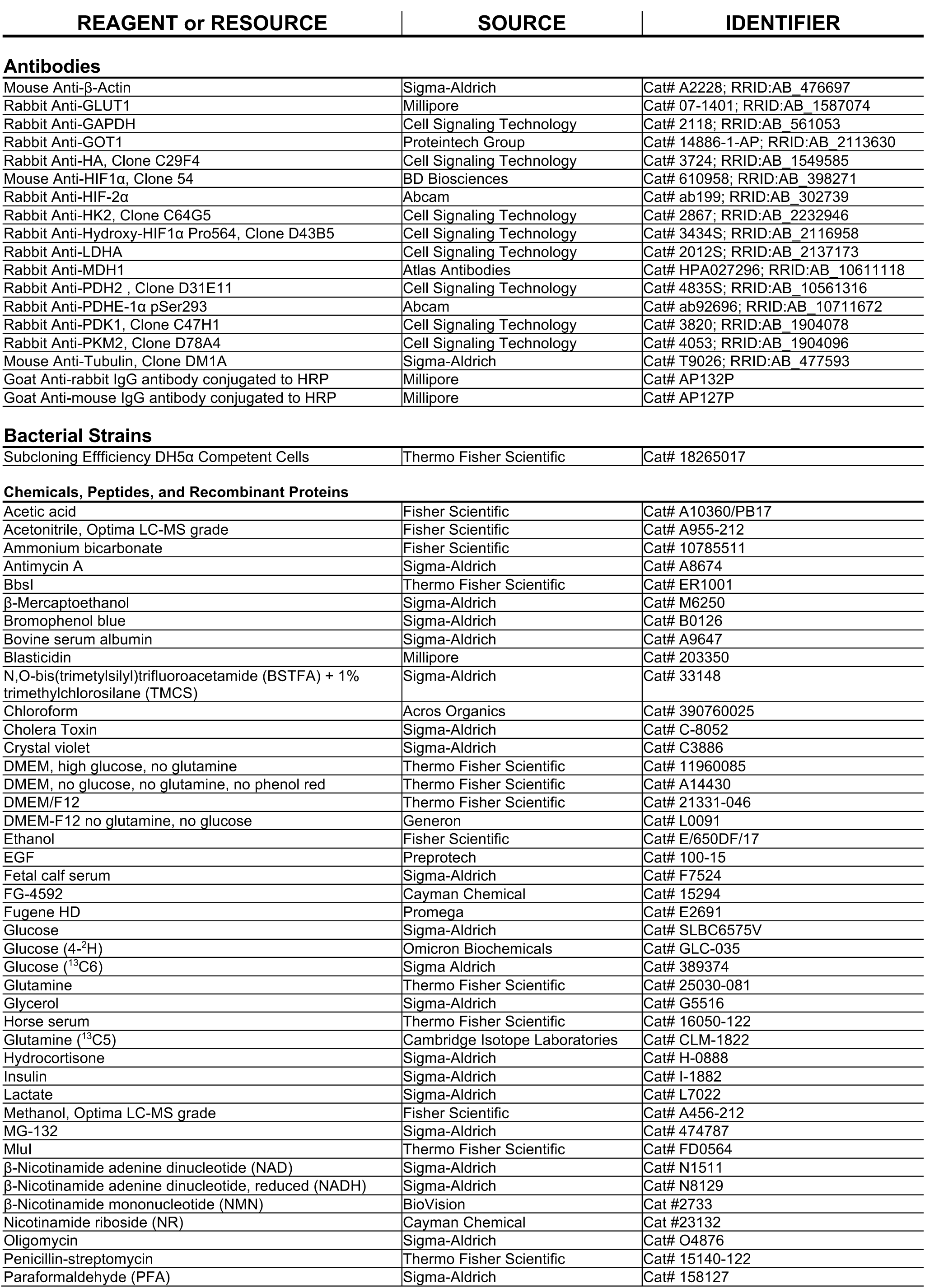
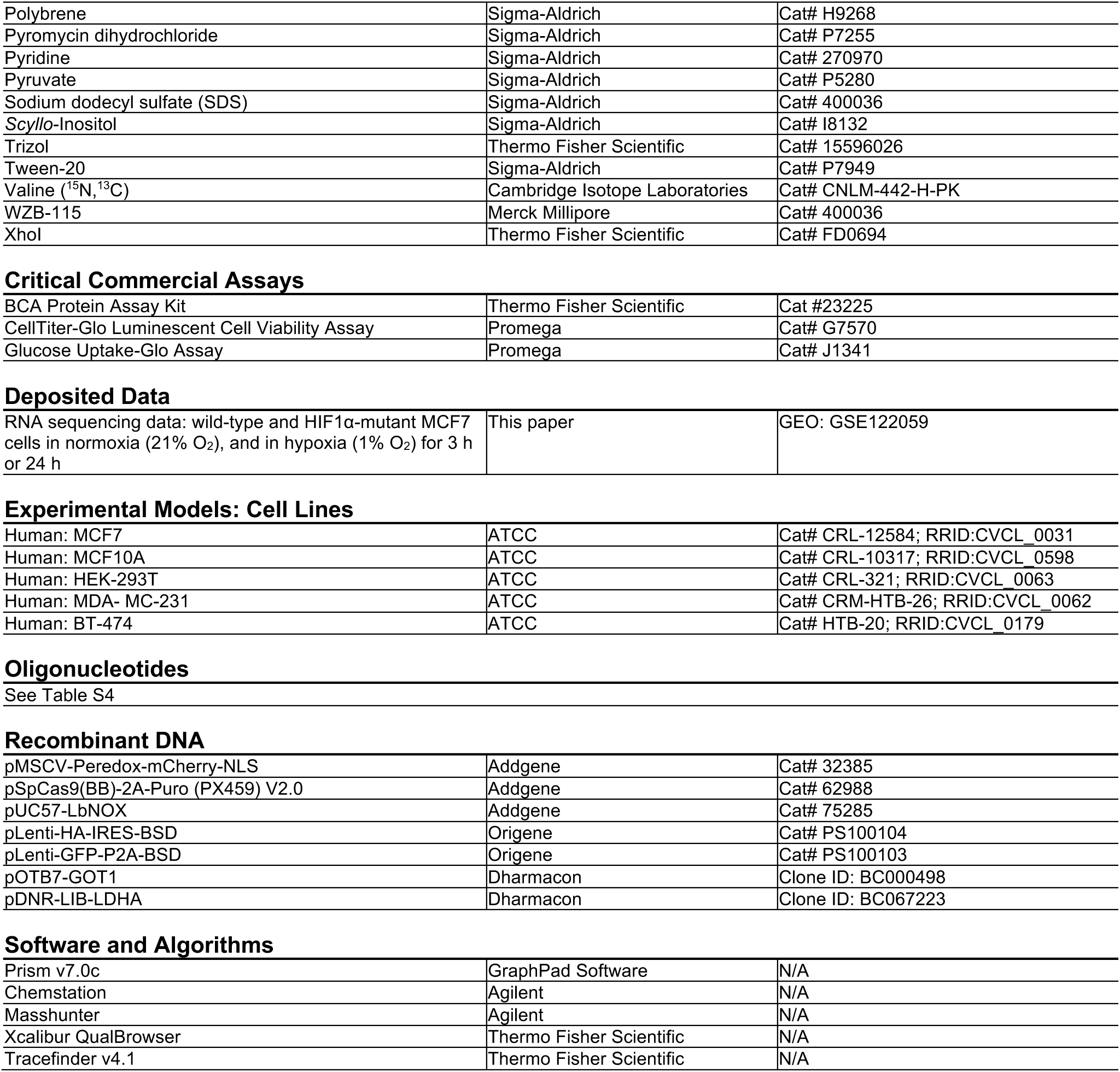

| REAGENT or RESOURCE | SOURCE | IDENTIFIER |
| --- | --- | --- |
| **Antibodies** |  |  |
| Mouse Anti-β-Actin | Sigma-Aldrich | Cat# A2228; RRID:AB_476697 |
| Rabbit Anti-GLUT1 | Millipore | Cat# 07-1401; RRID:AB_1587074 |
| Rabbit Anti-GAPDH | Cell Signaling Technology | Cat# 2118; RRID:AB_561053 |
| Rabbit Anti-GOT1 | Proteintech Group | Cat# 14886-1-AP; RRID:AB_2113630 |
| Rabbit Anti-HA, Clone C29F4 | Cell Signaling Technology | Cat# 3724; RRID:AB_1549585 |
| Mouse Anti-HIF1α, Clone 54 | BD Biosciences | Cat# 610958; RRID:AB_398271 |
| Rabbit Anti-HIF-2α | Abcam | Cat# ab199; RRID:AB_302739 |
| Rabbit Anti-HK2, Clone C64G5 | Cell Signaling Technology | Cat# 2867; RRID:AB_2232946 |
| Rabbit Anti-Hydroxy-HIF1α Pro564, Clone D43B5 | Cell Signaling Technology | Cat# 3434S; RRID:AB_2116958 |
| Rabbit Anti-LDHA | Cell Signaling Technology | Cat# 2012S; RRID:AB_2137173 |
| Rabbit Anti-MDH1 | Atlas Antibodies | Cat# HPA027296; RRID:AB_10611118 |
| Rabbit Anti-PDH2, Clone D31E11 | Cell Signaling Technology | Cat# 4835S; RRID:AB_10561316 |
| Rabbit Anti-PDHE-1α pSer293 | Abcam | Cat# ab92696; RRID:AB_10711672 |
| Rabbit Anti-PDK1, Clone C47H1 | Cell Signaling Technology | Cat# 3820; RRID:AB_1904078 |
| Rabbit Anti-PKM2, Clone D78A4 | Cell Signaling Technology | Cat# 4053; RRID:AB_1904096 |
| Mouse Anti-Tubulin, Clone DM1A | Sigma-Aldrich | Cat# T9026; RRID:AB_477593 |
| Goat Anti-rabbit IgG antibody conjugated to HRP | Millipore | Cat# AP132P |
| Goat Anti-mouse IgG antibody conjugated to HRP | Millipore | Cat# AP127P |
| **Bacterial Strains** |  |  |
| Subcloning Efficiency DH5α Competent Cells | Thermo Fisher Scientific | Cat# 18265017 |
| **Chemicals, Peptides, and Recombinant Proteins** |  |  |
| Acetic acid | Fisher Scientific | Cat# A10360/PB17 |
| Acetonitrile, Optima LC-MS grade | Fisher Scientific | Cat# A955-212 |
| Ammonium bicarbonate | Fisher Scientific | Cat# 10785511 |
| Antimycin A | Sigma-Aldrich | Cat# A8674 |
| BbsI | Thermo Fisher Scientific | Cat# ER1001 |
| β-Mercaptoethanol | Sigma-Aldrich | Cat# M6250 |
| Bromophenol blue | Sigma-Aldrich | Cat# B0126 |
| Bovine serum albumin | Sigma-Aldrich | Cat# A9647 |
| Blasticidin | Millipore | Cat# 203350 |
| N,O-bis(trimethylsilyl)trifluoroacetamide (BSTFA) + 1% trimethylchlorosilane (TMCS) | Sigma-Aldrich | Cat# 33148 |
| Chloroform | Acros Organics | Cat# 390760025 |
| Cholera Toxin | Sigma-Aldrich | Cat# C-8052 |
| Crystal violet | Sigma-Aldrich | Cat# C3886 |
| DMEM, high glucose, no glutamine | Thermo Fisher Scientific | Cat# 11960085 |
| DMEM, no glucose, no glutamine, no phenol red | Thermo Fisher Scientific | Cat# A14430 |
| DMEM/F12 | Thermo Fisher Scientific | Cat# 21331-046 |
| DMEM-F12 no glutamine, no glucose | Generon | Cat# L0091 |
| Ethanol | Fisher Scientific | Cat# E/650DF/17 |
| EGF | Preprotech | Cat# 100-15 |
| Fetal calf serum | Sigma-Aldrich | Cat# F7524 |
| FG-4592 | Cayman Chemical | Cat# 15294 |
| Fugene HD | Promega | Cat# E2691 |
| Glucose | Sigma-Aldrich | Cat# SLBC6575V |
| Glucose (4-$^2$H) | Omicron Biochemicals | Cat# GLC-035 |
| Glucose ($^{13}$C6) | Sigma Aldrich | Cat# 389374 |
| Glutamine | Thermo Fisher Scientific | Cat# 25030-081 |
| Glycerol | Sigma-Aldrich | Cat# G5516 |
| Horse serum | Thermo Fisher Scientific | Cat# 16050-122 |
| Glutamine ($^{13}$C5) | Cambridge Isotope Laboratories | Cat# CLM-1822 |
| Hydrocortisone | Sigma-Aldrich | Cat# H-0888 |
| Insulin | Sigma-Aldrich | Cat# I-1882 |
| Lactate | Sigma-Aldrich | Cat# L7022 |
| Methanol, Optima LC-MS grade | Fisher Scientific | Cat# A456-212 |
| MG-132 | Sigma-Aldrich | Cat# 474787 |
| MluI | Thermo Fisher Scientific | Cat# FD0564 |
| β-Nicotinamide adenine dinucleotide (NAD) | Sigma-Aldrich | Cat# N1511 |
| β-Nicotinamide adenine dinucleotide, reduced (NADH) | Sigma-Aldrich | Cat# N8129 |
| β-Nicotinamide mononucleotide (NMN) | BioVision | Cat #2733 |
| Nicotinamide riboside (NR) | Cayman Chemical | Cat #23132 |
| Oligomycin | Sigma-Aldrich | Cat# O4876 |
| Penicillin-streptomycin | Thermo Fisher Scientific | Cat# 15140-122 |
| Paraformaldehyde (PFA) | Sigma-Aldrich | Cat# 158127 |
| Polybrene | Sigma-Aldrich | Cat# H9268 |
| Pyromycin dihydrochloride | Sigma-Aldrich | Cat# P7255 |
| Pyridine | Sigma-Aldrich | Cat# 270970 |
| Pyruvate | Sigma-Aldrich | Cat# P5280 |
| Sodium dodecyl sulfate (SDS) | Sigma-Aldrich | Cat# 400036 |
| Scyllo-Inositol | Sigma-Aldrich | Cat# I8132 |
| Trizol | Thermo Fisher Scientific | Cat# 15596026 |
| Tween-20 | Sigma-Aldrich | Cat# P7949 |
| Valine (<sup>15</sup>N, <sup>13</sup>C) | Cambridge Isotope Laboratories | Cat# CNLM-442-H-PK |
| WZB-115 | Merck Millipore | Cat# 400036 |
| Xhol | Thermo Fisher Scientific | Cat# FD0694 |
| Critical Commercial Assays |  |  |
| BCA Protein Assay Kit | Thermo Fisher Scientific | Cat #23225 |
| CellTiter-Glo Luminescent Cell Viability Assay | Promega | Cat# G7570 |
| Glucose Uptake-Glo Assay | Promega | Cat# J1341 |
| Deposited Data |  |  |
| RNA sequencing data: wild-type and HIF1α-mutant MCF7 cells in normoxia (21% O<sub>2</sub>), and in hypoxia (1% O<sub>2</sub>) for 3 h or 24 h | This paper | GEO: GSE122059 |
| Experimental Models: Cell Lines |  |  |
| Human: MCF7 | ATCC | Cat# CRL-12584; RRID:CVCL_0031 |
| Human: MCF10A | ATCC | Cat# CRL-10317; RRID:CVCL_0598 |
| Human: HEK-293T | ATCC | Cat# CRL-321; RRID:CVCL_0063 |
| Human: MDA- MC-231 | ATCC | Cat# CRM-HTB-26; RRID:CVCL_0062 |
| Human: BT-474 | ATCC | Cat# HTB-20; RRID:CVCL_0179 |
| Recombinant DNA |  |  |
| pMSCV-Peredox-mCherry-NLS | Addgene | Cat# 32385 |
| pSpCas9(BB)-2A-Puro (PX459) V2.0 | Addgene | Cat# 62988 |
| pUC57-LbNOX | Addgene | Cat# 75285 |
| pLenti-HA-IRES-BSD | Origene | Cat# PS100104 |
| pLenti-GFP-P2A-BSD | Origene | Cat# PS100103 |
| pOTB7-GOT1 | Dharmacon | Clone ID: BC000498 |
| pDNR-LIB-LDHA | Dharmacon | Clone ID: BC067223 |
| Software and Algorithms |  |  |
| Prism v7.0c | GraphPad Software | N/A |
| Chemstation | Agilent | N/A |
| Masshunter | Agilent | N/A |
| Xcalibur QualBrowser | Thermo Fisher Scientific | N/A |
| Tracefinder v4.1 | Thermo Fisher Scientific | N/A |

### Cell lines and cell culture

Cell lines (MCF7, female, ATCC Cat# CRL-12584, RRID:CVCL_0031; MCF10A, female, ATCC Cat# CRL-10317, RRID:CVCL_0598; HEK-293T, female, ATCC Cat# CRL-3216, RRID:CVCL_0063; MDA-MB-231, female, ATCC Cat# CRM-HTB-26, RRID:CVCL_0062; BT-474, female, ATCC Cat# HTB-20, RRID:CVCL_0179) were obtained from the American Type Culture Collection (ATCC, Manassas, VA, USA). All cell lines were tested mycoplasma-free and cell identity was confirmed by short tandem repeat (STR) profiling by The Francis Crick Institute Cell Services Science Technology Platform. Cells (except MCF10A) were cultured in high-glucose DMEM (Gibco, Cat# 11960-085) supplemented with 10% fetal calf serum (FCS), 2 mM L-glutamine and 100 U/mL penicillin/streptomycin in a humidified incubator at 37°C, 5% CO_2_. MCF10A cells were cultured in DMEM/F12 (Gibco, Cat# 21331046), supplemented with 5% horse serum, 2 mM L-glutamine, 20 ng/ml EGF (PreproTech), 0.5 μg/ml hydrocortisone, 100 ng/ml cholera toxin, 10 μg/ml insulin and penicillin-streptomycin ^119^.

Prior to experiments, cells (except MCF10A) were seeded in glucose-free DMEM (Gibco, A1443001), supplemented with 5.5 mM glucose, 10% dialysed FCS (MWCO 3500), 2 mM L-glutamine and penicillin-streptomycin. MCF10A cells were seeded in DMEM/F12 (Generon Cat# L0091) supplemented with 5.5 mM glucose, 5% dialysed horse serum (MWCO 3500), 2 mM L-glutamine, 20 ng/ml EGF (PreproTech), 0.5 μg/ml hydrocortisone, 100 ng/ml cholera toxin, 10 μg/ml insulin and penicillin-streptomycin.

### DNA plasmids and cloning

pMSCV-Peredox-mCherry-NLS (Addgene plasmid # 32385) was a gift from Gary Yellen ^76^; pSpCas9(BB)-2A-Puro (PX459) V2.0 (Addgene plasmid # 62988) was a gift from Feng Zhang_120_. Plasmids pLenti-HA-IRES-BSD (PS100104) and pLenti-GFP-P2A-BSD (PS100103) were from Origene. pUC57-*Lb*NOX was a gift from Vamsi Mootha (Addgene plasmid # 75285) and used to subclone *Lb*NOX into the MluI and XhoI sites of pLenti-HA-IRES-BSD. The plasmids containing the cDNA of human GOT1 (Clone ID: BC000498) and human LDHA (Clone ID: BC067223) were obtained from Dharmacon. The cDNAs were amplified (GOT1 forward primer: cgc**acgcgt***acc*ATGGCACCTCCGTCAGTC, GOT1 reverse primer: gcg**ctcgag**CTGGATTTTGGTGACTGCTTC; LDHA forward primer: cgc**acgcgt***acc*ATGGCAACTCTAAAGGATCAG; LDHA reverse primer: gcg**ctcgag**AAATTGCAGCTCCTTTTGGATC*; lowercase italics*=Kozak sequence; **lowercase bold**=restriction enzyme cleavage site; UPPERCASE=insert) and cloned into pLenti-HA-IRES-BSD using the MluI and XhoI sites for lentivirus production. Results from experiments where this construct was used were compared to a control cell line transduced with virus encoding empty pLenti-GFP-P2A-BSD vector.

### Hypoxia treatments

Hypoxia treatments were performed in a Baker Ruskinn InVivo_2_ 400 hypoxic workstation at 1% O_2_, 5% CO_2_, 37°C and 70% humidity. Before each experiment, the media of cells were exchanged with media that had been pre-equilibrated at 1% O_2_ overnight as soon as cells were transferred to the hypoxia workstation. For hypoxia treatments prior to metabolomics experiments see *Stable isotope labelling and metabolite extraction*.

### Virus production and cell transduction

Retroviruses were produced in HEK-293T cells by co-transfecting pMSCV-Peredox-mCherry-NLS and a plasmid containing the amphotropic receptor gene (pHCMV-AmphoEnv) using FuGENE HD Transfection Reagent (Promega). Viral supernatants were harvested 48 h and 72 h after transfection, filtered and supplemented with 4 μg/mL polybrene. Viral supernatants were added to target cells for 6-8 h and cells were allowed to recover for 24 h prior to selection with 1 μg/mL puromycin for at least 5 days.

Lentiviral production and cell transduction were performed as for retroviruses, except that HEK-293T cells were transfected with pLenti-based vectors together with pMD2.G (VSV-G), pMDLg/pRRE (GAG/POL) and pRSV-Rev. Cells were selected with 5 μg/mL blasticidin for at least 5 days.

### Generation of knockout cell lines using CRISPR/Cas9

CRISPR-Cas9 expression constructs were designed and cloned into pSpCas9(BB)-2A-Puro as previously described ^120^ with minor modifications detailed below. CRISPR guide sequences (sgRNAs) were designed using the MIT CRISPR Design Tool (crispr.mit.edu): (HIF1α_forward: caccgTTCTTTACTTCGCCGAGATC, HIF1α _reverse: aaacGATCTCGGCGAAGTAAAGAAc, GOT1_forward: caccgAGTCTTTGCCGAGGTTCCGC,

GOT1_reverse: aaacGCGGAACCTCGGCAAAGACTc; GOT2_forward: caccgTGGAAGGCGGCGGCGATCCC; GOT2_reverse: aaacGGGATCGCCGCCGCCTTCCAc LDHA_forward: caccGGCTGGGGCACGTCAGCAAG, LDHA_reverse: aaacCTTGCTGACGTGCCCCAGCC). Corresponding guide oligonucleotides were mixed, phosphorylated using T4 Polynucleotide Kinase (New England Biolabs) and annealed in a thermocycler using the following programme: 37°C for 30 min, 95°C for 5 min, decrease temperature to 25°C at 0.1°C/min. The empty Cas9 expression plasmid was linearised using BbsI, ligated to annealed oligonucleotides and transformed into DH5α *E. coli*. Colonies were tested for successful insertion by colony PCR using the forward primer AATTTCTTGGGTAGTTTGCAGTTTT and the reverse guide oligonucleotide, with an expected band at 150bp.

MCF7 cells (70-90% confluency) were transfected with the CRISPR-Cas9 expression constructs using FuGENE HD Transfection Reagent (Promega), according to the manufacturer’s instructions. The day after transfection 1 μg/mL puromycin was added to the medium for 72 h before cells were seeded at limiting dilutions to obtain monoclonal colonies (500-1000 cells per 15 cm cell culture dish). After 2 weeks colonies (>100 cells) were isolated and expanded until they could be tested for loss of the target protein by western blot.

### Validation of HIF1α knockout in MCF7 cells

To verify the sequence of mutated HIF1α alleled in HIF1α-mutant MCF7 cells, genomic DNA was extracted using the NucleoSpin Blood kit (Macherey-Nagel), according to the manufacturer’s instructions. Exon 2 of the HIF1α gene, which had been targeted using CRISPR/Cas9, was amplified (forward primer: TTCCATCTCGTGTTTTTCTTGTTGT, reverse primer: CAAAACATTGCGACCACCTTCT) and PCR products were resolved on a 2% agarose gel. Bands around the expected size (317 bp) were purified and ligated into the pCR4Blunt-TOPO vector using the Zero Blunt TOPO PCR Kit for Sequencing (Thermo Fisher Scientific). Ligation products were transformed into One Shot TOP10 Chemically Competent *E. coli* cells (Thermo Fisher Scientific). *E. coli* cells were plates on LB plates containing kanamycin with X-gal (20 μl per plate, 8% w/v in dimethylformamide) and plasmid DNA was amplified and sequenced from 10 blue *E. coli* colonies (M13 forward primer: TGTAAAACGACGGCCAGT).

### Cell mass accumulation assay (crystal violet staining)

Cells were seeded in 24-well plates (50,000 cells/well) and, after the indicated treatments, cells were washed twice with PBS, fixed with 4% PFA, pH 7.4 (15 min, room temperature), washed with PBS and stained with 0.1% crystal violet in 20% methanol. After staining, cells were washed twice with distilled water (10 min each) and dried. After re-solubilisation in 10% acetic acid, absorbance at 595 nm was measured using a Tecan infinite M1000 Pro plate reader.

### Continuous cell proliferation assay

Cells were seeded the day before the experiment in 96-well plates (black, transparent bottom, Corning, #3606; 9,000-12,000 cells/well). After addition of treatments and/or changing the oxygen concentration in the incubator to 1% O_2_, cell confluence was monitored using an IncuCyte S3 (Essen Bioscience) by taking phase-contrast images using a 10x objective.

### End-point cell proliferation assay

Cells were seeded the day before the experiment in 96-well plates (black, transparent bottom, Corning, #3606; 9,000-12,000 cells/well). 24 hours after addition of treatments and/or changing the oxygen concentration in the incubator to 1% O_2_ cells were fixed by adding an equal volume (100 μl) of 8% paraformaldehyde, pH7.4 and incubated for 10 minutes at room temperature. Cells were washed with PBS and stored in PBS at 4°C until further processing. Cells were stained with DAPI (4’,6-diamidino-2-phenylindole, 1 μg/ml) for 1 hour at room temperature, washed twice with PBS and nuclei counting was performed using an Acumen Explorer eX3 laser scanning microplate cytometer (TTP Labtech).

### Cell lysis and western blotting

Cells were washed twice with PBS and lysed in Laemmli buffer (50 mM Tris-HCl pH 6.8, 1% SDS, 10% glycerol) and stored at −20°C until further use. Lysed samples were sonicated (2− 10 s), protein concentration was measured and samples were boiled at 95 °C for 5 min after addition of 5% β-mercaptoethanol and bromophenol blue. Samples (20 μg of protein) were resolved by SDS-PAGE and proteins were transferred to nitrocellulose membranes by electroblotting. Membranes were blocked with 5% milk in TBS-T (50 mM Tris-HCl pH 7.5, 150 mM NaCl, 0.05% Tween-20) for 1 h at room temperature and incubated with the primary antibody overnight at 4°C. Membranes were washed three times with TBS-T and incubated with secondary antibody conjugated to horseradish peroxidase (1:2000) in 5% milk TBS-T for 1 h at room temperature. Antibodies were visualised by chemiluminescence and imaged using medical X-ray film developed in an AGFA Curix 60 processor (Figures 2A, 3A, 5A, S2A) or imaged using the Amersham Imagequant 600 RGB (Figures 3F, 6B-D, S4E, S6D-F). Primary antibodies used: mouse anti-β-actin antibody (Sigma-Aldrich Cat# A2228, RRID:AB_476697), 1:2000 in 5% BSA/TBS-T; rabbit anti-GAPDH (Cell Signaling Technology Cat# 2118, RRID:AB_561053), 1:1000 in 5% BSA /TBS-T; rabbit anti-GLUT1 (Millipore Cat# 07-1401, RRID:AB_1587074), 1:5000 in 5% milk /TBS-T; rabbit anti-GOT1 (Proteintech Group Cat# 14886-1-AP, RRID:AB_2113630), 1:250 in 5% BSA /TBS-T; rabbit anti-HA (Clone C29F4, Cell Signaling Technology Cat# 3724, RRID:AB_1549585), 1:1000 in 5% BSA /TBS-T; mouse anti-HIF1α (Clone 54, BD Biosciences Cat# 610958, RRID:AB_398271), 1:250 in 5% milk/TBS-T; rabbit anti-HIF-2α (Abcam Cat# ab199, RRID:AB_302739), 1:1000 in 5% BSA/TBS-T; rabbit anti-HK2 (Clone C64G5, Cell Signaling Technology Cat# 2867, RRID:AB_2232946), 1:1000 in 5% BSA /TBS-T; rabbit anti-Hydroxy-HIF1α Pro564 (Clone D43B5, Cell Signaling Technology Cat# 3434S, RRID:AB_2116958), 1:1000 in 5% BSA /TBS-T; rabbit anti-LDHA (Cell Signaling Technology Cat# 2012S, RRID:AB_2137173), 1:1000 in 5% BSA /TBS-T; rabbit anti-MDH1 (Atlas Antibodies Cat# HPA027296, RRID:AB_10611118), 1:500 in 5% BSA /TBS-T; rabbit anti-PDH2 (clone D31E11, Cell Signaling Technology Cat# 4835S, RRID:AB_10561316), 1:500 in 5% BSA /TBS-T; rabbit anti-PDHE-1α pSer293 (Abcam Cat# ab92696, RRID:AB_10711672), 1:500 in 5% BSA /TBS-T; rabbit anti-PDK1 (Clone C47H1, Cell Signaling Technology Cat# 3820, RRID:AB_1904078), 1:100 in 5% BSA /TBS-T; rabbit anti-PKM2 (Clone D78A4, Cell Signaling Technology Cat# 4053, RRID:AB_1904096), 1:1000 in 5% BSA /TBS-T; mouse anti-Tubulin (Clone DM1A, Sigma-Aldrich Cat# T9026, RRID:AB_477593), 1:2000 in 5% milk/TBS-T; Secondary antibodies (Millipore): Goat anti-rabbit IgG antibody conjugated to HRP, goat anti-mouse IgG antibody conjugated to HRP.

### Transcriptional profiling by RNA sequencing

Cells were seeded 48 h before harvest (1.5 x 10^6^ cells per 6 cm plate) and after indicated treatments (3 h or 24 h at 1% O_2_, or medium change 3 h before harvest in control cells), cells were washed three times with PBS and lysed in 1 ml TRIzol Reagent (Thermo Fisher Scientific). Chloroform (0.2 ml, Acros Organics) was added to lysates and after shaking for 5 seconds samples were centrifuged for 18 min at 10,000 x *g*. The upper, aqueous phase was mixed with an equal volume of 100% ethanol (500 μl) and RNA was purified using the RNeasy Kit (Qiagen), according to the manufacturer’s instructions. DNase treatment was omitted since initial tests showed no contamination by genomic DNA. After RNA quantification and quality control (NanoDrop, Qubit and Agilent 2100 Bioanalyzer), libraries were prepared using the TruSeq RNA Library Prep Kit v2 (Illumina) or KAPA mRNA HyperPrep Kit (Kapa Biosystems). mRNA sequencing was performed on a Illumina HiSeq 2500 instrument (paired-end or single-end reads, 25 million reads total).

### ATP quantification assay

Cells were seeded in 96-well plates (15,000 cells per) 24-48 h before measurement. After treatment with hypoxia for the indicated lengths of time, or the indicated compounds for 3 h, ATP-dependent luciferase luminescence was measured using the CellTiterGlo kit (Promega, Cat# G7570) according to the manufacturer’s instructions. Luminescence counts were normalised to cell number.

### Glucose uptake

Cells were seeded in 96-well plates (15,000 cells per well) 48 h prior to measurement. 2DG uptake was measured using the Glucose Uptake-Glo Assay kit (Promega) after indicated hypoxia treatments (or medium change 3 h before the experiment for control cells) and incubation with 1 mM 2DG for 10 min.

For measurements under hypoxic conditions, 2DG was added to cells in the hypoxia workstation, the plates were sealed with several layers of parafilm, transferred to ambient atmosphere and incubated for 10 min. After adding stop and neutralisation reagents, samples were transferred to a white 96-well plate and incubated for 1 h before luminescence was measured using a Tecan infinite M1000 Pro plate reader. Luminescence counts were normalised to cell number. Wells containing cells without 2DG as well as cells treated with the GLUT1 inhibitor WZB-115 (50 μM for 15 min, #400036, Merck) were used as negative controls (not shown).

### Mitochondrial oxygen consumption

Cells were seeded 48 h before the experiment and, after the indicated treatments, cells were trypsinised, centrifuged (1500 rpm, 3 min, room temperature) and resuspended at 5-7.5 x10^5^ cells/mL. Hypoxia-treated cells were processed in the hypoxic workstation and re-suspended in medium pre-equilibrated in a hypoxic atmosphere (as in **Hypoxia treatments**). Respiration was measured using an O2k oxygraph (Oroboros instruments) as previously described ^121,122^ in sealed chambers to prevent re-oxygenation of hypoxia-treated cells. Residual oxygen consumption (ROX) was measured after the addition of the complex III inhibitor antimycin A (2.5 μM) and was subtracted from basal oxygen consumption. Oxygen consumption was corrected for cell number.

### Imaging of cytoplasmic NADH with Peredox

MCF7 cells stably expressing Peredox ^76^ were seeded in glass-bottom 24-well plates the day before the experiment (100,000 cells/well). One hour before the experiment, the medium was replaced with imaging buffer (10 mM HEPES pH 7.4, 140 mM NaCl, 1 mM CaCl_2_, 1 mM MgCl_2_, 5.4 mM KCl) containing 5.5 mM glucose and 2 mM L-glutamine to acquire baseline images. After image acquisition, cells were incubated with imaging buffer containing 10 mM lactate or 10 mM pyruvate for 5 min before imaging. Cells were kept at 37°C throughout the experiment and washed with warm imaging buffer between treatments.

Fluorescence images were acquired using an AxioObserver Z1 microscope (Zeiss) and Plan-Apochromat 20x/0.8 M27 objective (picture size 512−512 pixels, 0.6x zoom, 600.3 μm pinhole. T-sapphire was excited at 800 nm (10% laser power, gain 800) using a Mai Tai DeepSee laser (Spectra-Physics) and emission was recorded at 525 nm. Images were processed using Fiji software by converting pictures from vendor format to 8-bit tiff format, thresholding and identifying nuclei using the ‘Analyze particle’ function. Mean intensity per particle was used to calculate changes of T-sapphire fluorescence in individual nuclei from baseline to treatment with lactate or pyruvate.

### Stable isotope labelling and metabolite extraction

Metabolomics sample preparation, GC-MS data processing and analysis were performed as described in ^123^. In brief, cells were seeded in 6-well plates (0.35-0.5×10^6^ cells per well, 4-5 replicates per condition) 24-48 h before harvest. One hour before the start of the experiment the medium was refreshed and was again changed to medium containing the isotopically labelled nutrient (5.5 mM [U-^13^C]-glucose, 5.5 mM [4-^2^H]-glucose or 2 mM [U-^13^C]-glutamine) at the start of the experiment. Cells were washed twice with PBS, immediately quenched with liquid nitrogen and kept on dry ice until extraction. Metabolites were extracted by scraping cells in 500 μl methanol, followed by washing the plate with 250 μl methanol and 250 μl water containing the polar internal standard *scyllo*-inositol (1 nmol per sample). Fractions were combined with 250 μl chloroform containing the apolar internal standard [1-_13_C]-lauric acid (C12:0, 40 nmol per sample). Extracts were vortexed, sonicated for 3 x 8 min and incubated at 4°C overnight. Precipitate was removed by centrifugation (10 min, 18,000 x *g*, 4°C) and phases were separated by adding 500 μl water (resulting in 1:3:3 (v/v) chloroform/methanol/water) and centrifugation (5 min, 18,000 x *g*, 4°C). In parallel, cells from 3 wells per experimental condition were trypsinised and counted using a Nexcelcom Bioscience Cellometer Auto T4 for subsequent normalisation of data to cell number.

The protocol used to extract NAD^+^ and NADH was adapted from ^78^. Briefly, cells were scraped in 250 μl acetonitrile:methanol:20mM ammonium bicarbonate pH 9.0 (2:2:1 v/v) containing 5 μM ^15^N^13^C-valine as internal standard, followed by sonication for 3 x 8 min and incubation at 4°C for 1 h. Precipitate was removed by centrifugation, samples were transferred to glass vials and immediately analysed by LC-MS.

### Gas chromatography-mass spectrometry (GC-MS)

For GC-MS analysis of intracellular metabolites, aqueous phases of cell extracts were transferred to glass vial inserts. For analysis of metabolites in media, 5 μl of media samples were transferred to glass vial inserts and spiked with 1 nmol *scyllo*-inositol. Samples were dried in a centrifugal evaporator and washed twice with 40 μl methanol followed by drying. Samples were methoxymated (20 μl of 20 mg/mL methoxyamine in pyridine, at room temperature overnight) and derivatised with 20 μl of *N*,*O*-bis(trimetylsilyl)trifluoroacetamide (BSTFA) + 1% trimethylchlorosilane (TMCS) for at least 1 h. GC-MS analysis of metabolites was performed using Agilent 7890B-5977A and 7890A-5975C systems in splitless injection mode (1 μl of sample, injection temperature 270°C) with a DB-5MS DuraGuard column, helium as carrier gas and electron impact ionization. Oven temperature was initially 70 °C (2 min), followed by a temperature increase to 295 °C at 12.5 °C/min and subsequently to 320 °C at 25 °C/min (held for 3 min). Chemstation and MassHunter Workstation software (B.06.00 SP01, Agilent Technologies) was used for metabolite identification and quantification by comparison to the retention times, mass spectra, and responses of known amounts of authentic standards. Internal standards were used to correct for sample losses during phase separation and metabolite abundances were normalised to cell number. See **Table S2** for fragment ions used for metabolite quantification by GC-MS.

### Liquid chromatography-mass spectrometry (LC-MS)

Aqueous phases were transferred to glass vial inserts and dried in a centrifugal evaporator. Dried metabolites were resuspended in 100 μl methanol:water (1:1 v/v). The LC-MS method was adapted from ^124^. LC-MS analysis was performed using a Dionex UltiMate LC system (Thermo Scientific) with a ZIC-pHILIC column (150 mm x 4.6 mm, 5 μm particle, Merck Sequant). A 15 min elution gradient of 80% Solvent A (20 mM ammonium carbonate in Optima HPLC grade water, Sigma Aldrich) to 20% Solvent B (acetonitrile Optima HPLC grade, Sigma Aldrich) was used, followed by a 5 min wash of 95:5 Solvent A to Solvent B and 5 min re-equilibration. Other parameters were as follows: flow rate, 300 μL/min; column temperature, 25°C; injection volume, 10 μL; autosampler temperature, 4°C.

MS was performed in positive/negative polarity switching mode using a Q Exactive Orbitrap instrument (Thermo Scientific) with a HESI II (Heated electrospray ionization) probe. MS parameters were as follows: spray voltage 3.5 kV for positive mode and 3.2 kV for negative mode; probe temperature, 320°C; sheath gas, 30 arbitrary units; auxiliary gas, 5 arbitrary units; full scan range: 70 to 1050 m/*z* with settings of AGC target and resolution as ‘Balanced’ and ‘High’ (3×10^6^ and 70,000), respectively. Data were recorded using Xcalibur 3.0.63 software (Thermo Scientific). Mass calibration was performed for both ESI polarities before and lock-mass correction was applied to each analytical run using ubiquitous low-mass contaminants to enhance calibration stability. Pooled biological quality control (PBQC) samples were prepared by pooling equal volumes of each sample, and were analysed throughout the run to provide a measurement of the stability and performance of the instrument. PBQC samples were also analysed in parallel reaction monitoring (PRM) mode to confirm the identification of metabolites. PRM acquisition parameters: resolution 17,500, auto gain control target 2−10^5^, maximum isolation time 100 ms, isolation window m/z 0.4; collision energies were set individually in HCD (high-energy collisional dissociation) mode. Qualitative and quantitative analyses were performed using Xcalibur Qual Browser and Tracefinder 4.1 software (Thermo Scientific) according to the manufacturer’s workflows. See **Table S3** for fragment ions used for metabolite quantification by LC-MS.

### Quantification and statistical analysis

Statistical analyses were performed using R ^125^ or GraphPad Prism v7.0c. Comparisons were made using either two-tailed, unpaired t-tests, one-way or two-way ANOVA with Dunnett’s correction, Sidak’s correction or Tukey’s correction for multiple comparisons, as indicated in the respective figure legends. All error bars shown in graphs and measurement error in the text (±) represent standard deviation. Statistical errors were propagated where indicated in the figure legends. Significance levels are defined as follows: * p<0.05, ** p<0.01, *** p<0.001, **** p<0.0001. Curve-fitting was performed using GraphPad Prism v7.0c using a one-phase decay model with standard parameters. Western blot images were quantified using Fiji ImageJ v1.45 software.

GC-MS metabolomics data were analysed using Agilent Chemstation and MassHunter software, as well as in-house R scripts. Qualitative and quantitative analysis of LC-MS metabolomics data was performed using Xcalibur QualBrowser and Tracefinder 4.1 software (Thermo Fisher Scientific). Isotopic labelling data were corrected for natural isotope abundance using a script provided by Sean O’Callaghan (Bio21 Institute, The University of Melbourne). To plot heatmaps, data were expressed as log_2_-fold change relative to the mean of the control condition, averaged across all replicates per sample group and subsequently plotted using the RColorBewer v1.1-2 ^126^ and pheatmap R packages v1.0.8 ^127^. Z-scores were calculated by subtracting the mean of control condition from the measured value and dividing by the standard deviation of the control condition. All isotope labelling data are expressed as fraction of labelled molecules per metabolite, unless specified otherwise in the figure legends.

Analysis of RNA sequencing data was performed in the R environment and controlled by a GNU make pipeline. Transcript reads were aligned to the Ensembl GRCh37 genome using Tophat2 v2.1.1 ^128^. Aligned transcript reads were filtered for genes with at least 10 reads per gene in 5 or more samples. Between-sample normalisation was performed using the RUVSeq R package v1.10.0 ^129^ and differential expression between sample groups was evaluated using the DESeq2 package v1.16.1 ^130^. The edgeR package v3.18.1 ^131^ was used to calculate false discovery rates (FDR) and a cut-off of 1% was applied. Ensembl IDs were converted to gene names and entrez IDs using the AnnotationDBI v1.40.0 ^132^, EnsDb.Hsapiens.v79 v2.99.0, ensembldb v2.2.2 ^133^ and org.Hs.eg.db v3.5.0 ^134^ packages.

## DATA AND SOFTWARE AVAILABILITY

The computer code used in this study is available from the corresponding author upon reasonable request. The data that support the findings of this study are available from the corresponding author upon reasonable request. The RNA sequencing data discussed in this publication have been deposited in NCBI’s Gene Expression Omnibus and are accessible through GEO Series accession number GSE122059.

---

**Figure S1 related to Figure 1.**
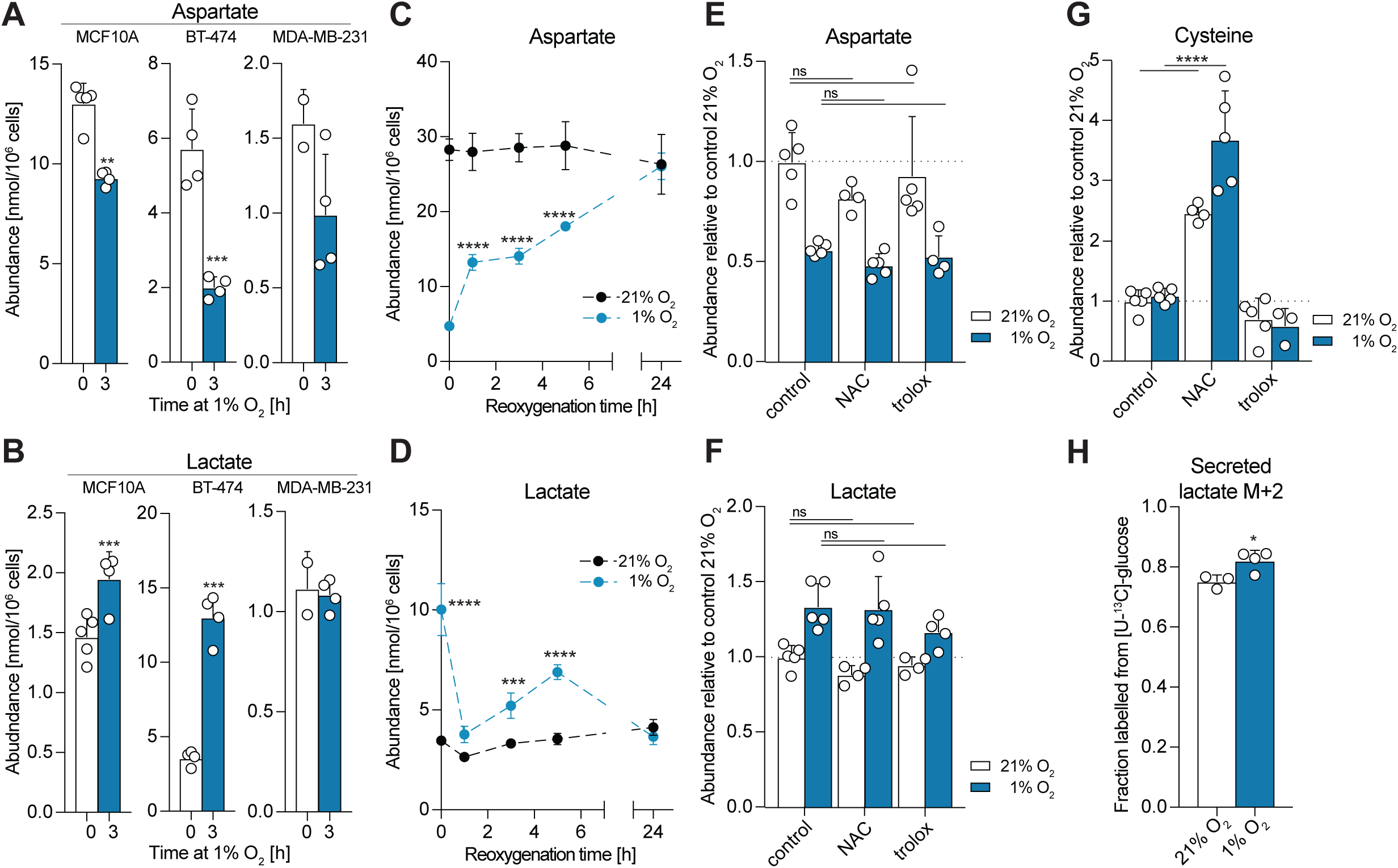
Increase in lactate and decrease in aspartate levels induced by exposure to 1% O_2_ occur in other cell lines, are reversible upon reoxygenation and cannot be prevented by antioxidant treatment. **A-B)** Intracellular abundances of lactate and aspartate in the indicated cell lines at 21% O_2_ and after 3 h at 1% O_2_ (n= 2-5 cultures per cell line and condition). Data are shown as mean ±SD and significance relative to 21% O_2_ was tested using two-tailed, unpaired t-tests. See also Figure 1C-D. **C-D)** Intracellular abundances of lactate and aspartate in MCF7 cells exposed to 1% O_2_ for 3 h, followed by reoxygenation at 21% O_2_ for the indicated lengths of time, compared to control cells in normoxia (n = 4 cultures per time point and condition). Data are shown as mean ± SD and significance relative to 21% O_2_ was tested using two-way ANOVA with Sidak’s multiple comparison test. **E-G)** Intracellular abundances of lactate, aspartate and cysteine in MCF7 cells incubated in 21% O_2_ or 1% O_2_ for 3 h, with and without antioxidant treatment (NAC, N-acetylcysteine: 1mM; Trolox: 50 μM; n = 5 cultures per and condition). Cysteine levels shown to confirm the effect of NAC on cells. Data are shown as mean ± SD and significance relative to 21% O_2_ was tested using two-way ANOVA with Dunnett’s multiple comparison test. **H)** Fraction of lactate M+2 labelled from [U-^13^C]-glucose in cell culture media from MCF7 cells incubated in 21% O_2_ or 1% O_2_ for 5 h with the isotopic tracer (n = 3 cultures per condition). The quantified ion (m/*z* 117) contains two lactate-derived carbons. Data are shown as mean ± SD and significance relative to 21% O_2_ was tested using a two-tailed, unpaired t-test.

**Figure S2 related to Figure 2.**
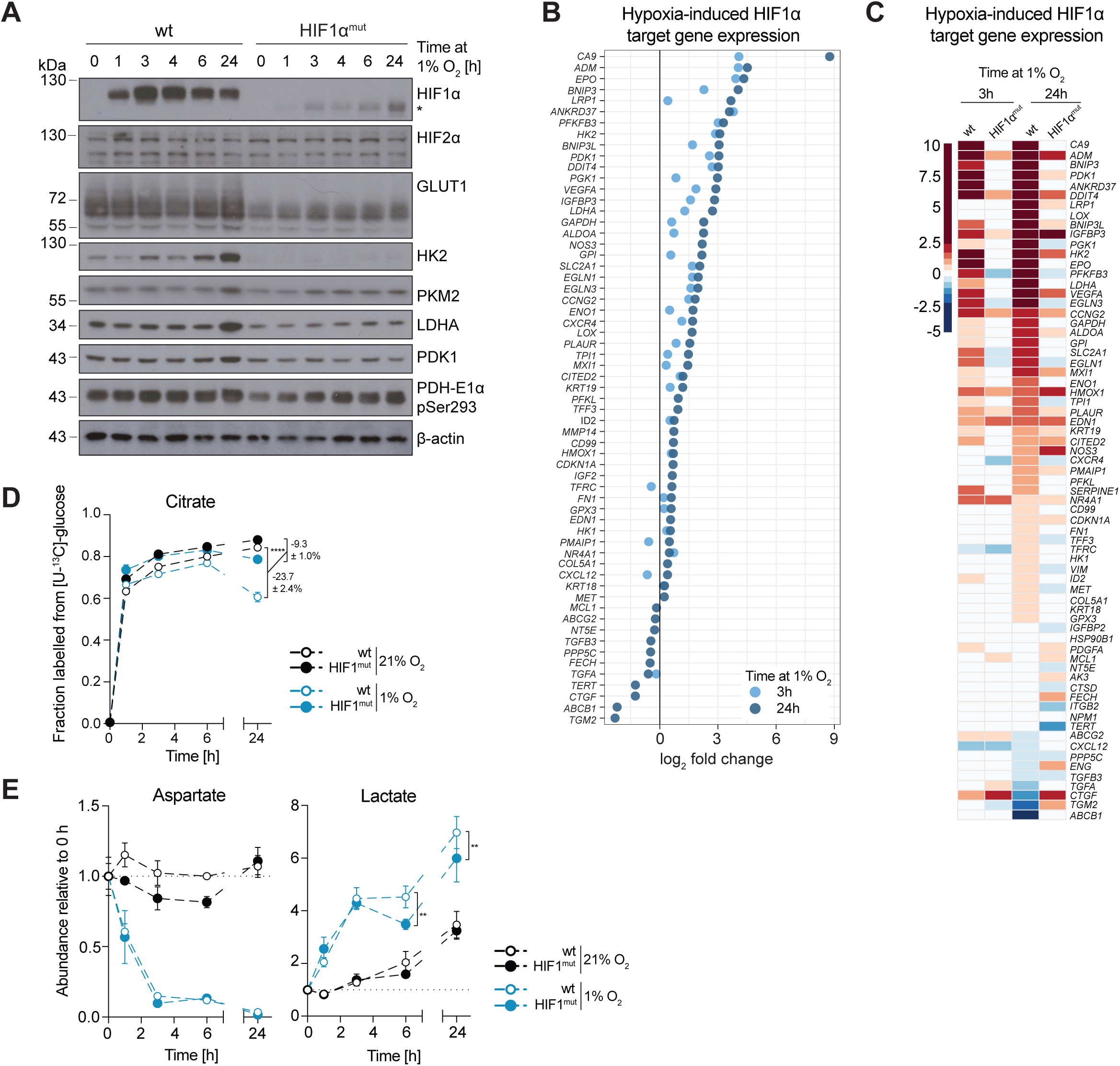
HIF1α target expression in breast cancer cell lines and metabolic response to hypoxia in HIF1α^mut^ MCF7 cells. **A)** Western blot to assess levels of HIF1α and a panel of HIF1α targets in the indicated cell lines incubated in 21% O_2_ or at 1% O_2_ for the indicated lengths of time. See also Figure 2A. **B)** Sequence alignment of mutant HIF1α alleles in HIF1α^mut^ MCF7 cells with exon 2 of human HIF1α cDNA (CCDS9753.1). **C)** Fraction of the indicated metabolite pools labelled from [U-^13^C]-glucose in wild-type (wt) and HIF1α^mut^ MCF7 cells as in Figure 2D (n = 4 cultures for each time point and condition). Data are shown as mean ± SD. Statistical errors were propagated to calculate the variance of change in isotopic labelling between normoxia and hypoxia for each cell line. Significance of this change between cell lines was then tested using two-way ANOVA with Sidak’s multiple comparison test. See also Figure 2D.

**Figure S3 related to Figure 3.**
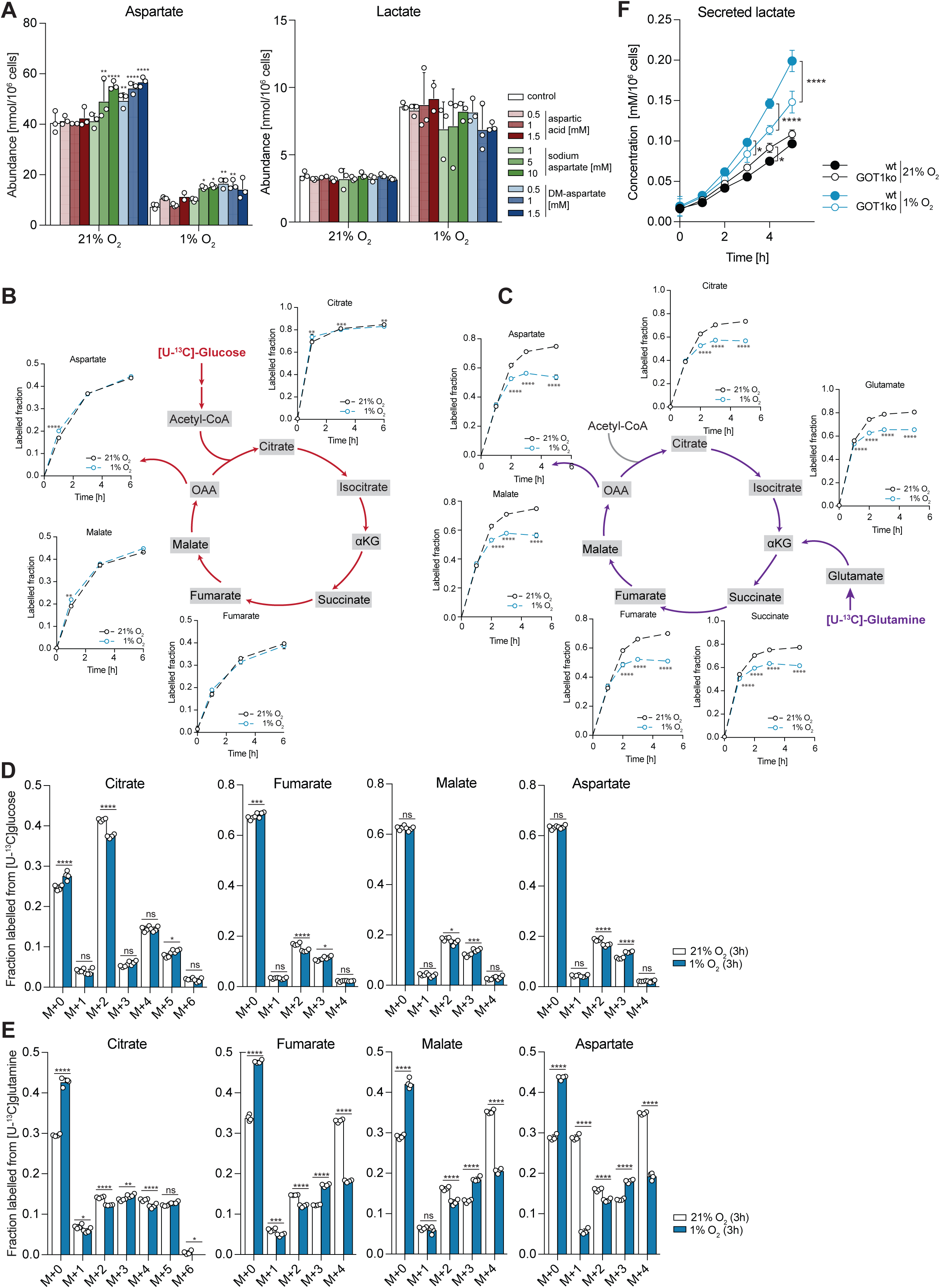
Exogenous aspartate does not change lactate levels in MCF7 cells – decreased labeling of aspartate from glutamine but not glucose – decreased level of secreted lactate in GOT1ko cells in hypoxia. **A)** Intracellular abundances of aspartate and lactate in MCF7 cells in 21% O_2_ and after 3 h in 1% O_2_. Media were supplemented with the indicated concentrations of aspartic acid, sodium aspartate or cell-permeable dimethyl-aspartate (DM-aspartate) for 3 h. Data are shown as mean ± SD (n = 2-3 cultures per condition) and significance relative to control cells in each condition (21% O_2_ or 1% O_2_) was tested using two-way ANOVA with Dunnett’s multiple comparison test. **B)** Fractions of the indicated metabolite pools labelled after incubation of MCF7 cells with [U-^13^C]-glucose in 21% O_2_ or 1% O_2_ for the lengths of time shown. Data are shown as mean ±SD (n = 4 cultures per condition and time point) and significance relative to 21% O_2_ was tested using two-way ANOVA with Sidak’s multiple comparison test. Data are from the same experiment as in Figure 2D and **S2C**, shown here for comparison with glutamine labeling in panel **C** below. **C)** As in **B** but using [U-^13^C]-glutamine. **D)** Fractions of indicated metabolite isotopologues labelled from [U-^13^C]-glucose after 3 hours incubation at 21% O_2_ or 1% O_2_. Data shown as mean ± SD (n = 4 cultures per condition, same experiment as in **B**) and significance relative not normoxia was tested using two-way ANOVA with Sidak’s multiple comparison test. **E)** Data shown as in **D** but using [U-^13^C]-glutamine (same experiment as in **C**). **F)** Lactate concentration in cell culture media of wild-type (wt) and GOT1ko MCF7 cells incubated in 21% O_2_ or 1% O_2_ for the indicated lengths of time (n = 3-4 cultures per time point and condition). Data are shown as mean ± SD and significance was tested using two-way ANOVA with Tukey’s multiple comparison test. See Figure 3E.

**Figure S4 related to Figure 4.**
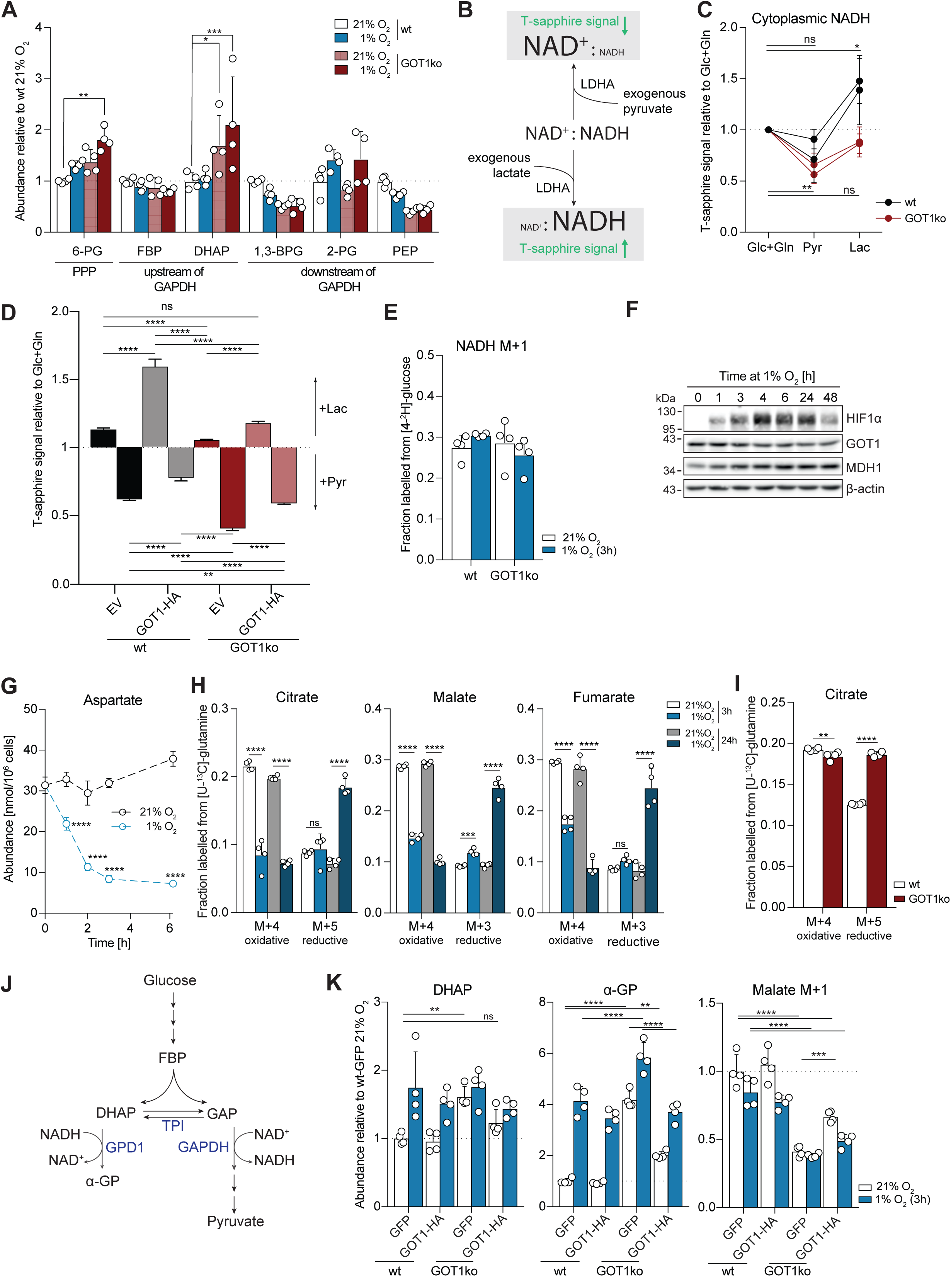
Aspartate metabolism, and reductive carboxylation in early hypoxia. **A)** Bar graph representation of the data in Figure 4A. Data shown as mean ± SD (n = 4 cultures per condition) and significance was tested using two-way ANOVA with Sidak’s multiple comparison test. **B)** Schematic illustrating the effect of exogenous lactate and pyruvate on the cytoplasmic NAD^+^/NADH ratio and Peredox T-sapphire fluorescence intensity. **C)** Peredox T-sapphire fluorescence signal intensity in wild-type (wt) and GOT1ko MCF7 cells in buffer containing 5.5 mM glucose and 2 mM glutamine (Glc+Gln) and after sequential incubation first with 10 mM pyruvate (Pyr) and then with 10 mM lactate (Lac) (reverse order of that used in Figure 4E). Signal was normalized per nucleus and is shown relative to the Glc+Gln condition. Data points represent mean ± SD of 2 independent replicates per cell line (n = 17-56 cells per replicate); significance relative to Glc+Gln was tested using one-way ANOVA with Dunnett’s multiple comparison test. See also Figure 4E. **D)** Peredox T-sapphire fluorescence signal intensity in wild-type (wt) and GOT1ko MCF7 cells stably expressing an empty vector (EV) or HA-tagged GOT1. Cells in buffer containing 5.5 mM glucose and 2 mM glutamine (Glc+Gln) were sequentially incubated first with 10 mM lactate (Lac) and then with 10 mM pyruvate (Pyr). Signal was normalized per nucleus and is shown relative to the Glc+Gln condition. Data points represent mean ± SEM (n = 64-654 cells per cell line, values from one representative of 3 independent experiments); significance relative to Glc+Gln was tested using Kruskal-Wallis test with multiple comparison. **E)** Fraction of NADH labelled from [4-^2^H]-glucose in the experiment shown in Figure 4G. Data shown as mean ± SD (n = 4 cultures per condition) and significance was tested using two-way ANOVA with Sidak’s multiple comparison test (not significant). **F)** Western blot to assess levels of GOT1 and MDH1 in wild-type (wt) and GOT1ko cells incubated in 1% O_2_ for the indicated lengths of time. **G)** Intracellular abundance of aspartate in the experiment shown in Figure 4H-J. Data shown as mean ± SD (n = 4 cultures per condition) and significance was tested using two-way ANOVA with Sidak’s multiple comparison test. **H)** Fractions of citrate, malate and fumarate labelled from [U-^13^C]-glutamine after incubation with the tracer in 21% O_2_ or 1% O_2_ for 3 or 24 h. Isotopologues produced via oxidative TCA (citrate M+4, malate M+4, fumarate M+4) and via reductive carboxylation (citrate M+5, malate M+3, fumarate M+3) are shown. Data shown as mean ± SD (n = 4 cultures per condition) and significance was tested using two-way ANOVA with Sidak’s multiple comparison test. **I)** Fraction of citrate labelled from [U-^13^C]-glutamine after incubation of wild-type (wt) and GOT1ko MCF7 cells with the tracer for 3 h. Data shown as mean ± SD (n = 4 cultures per condition) and significance was tested using two-way ANOVA with Sidak’s multiple comparison test. **J)** Schematic illustrating the fate of glucose carbons on either side of the triosephosphate isomerase (TPI) reaction. **K)** Intracellular abundance of dihydroxyacetone phosphate (DHAP), α-glycerophosphate (α-GP) and malate M+1 (labelled from [4-^2^H]-glucose after incubation with the tracer for 3 h) in wild-type (wt) and GOT1ko MCF7 cells stably expressing GOT1-HA or GFP after 3 h at 21% O_2_ or 1% O_2_. Data shown as mean ±SD (n = 4 cultures per condition) and significance was tested using two-way ANOVA with Sidak’s multiple comparison test (DHAP) or Tukey’s multiple comparison test (α-GP, Malate M+1).

**Figure S5 related to Figure 5.**
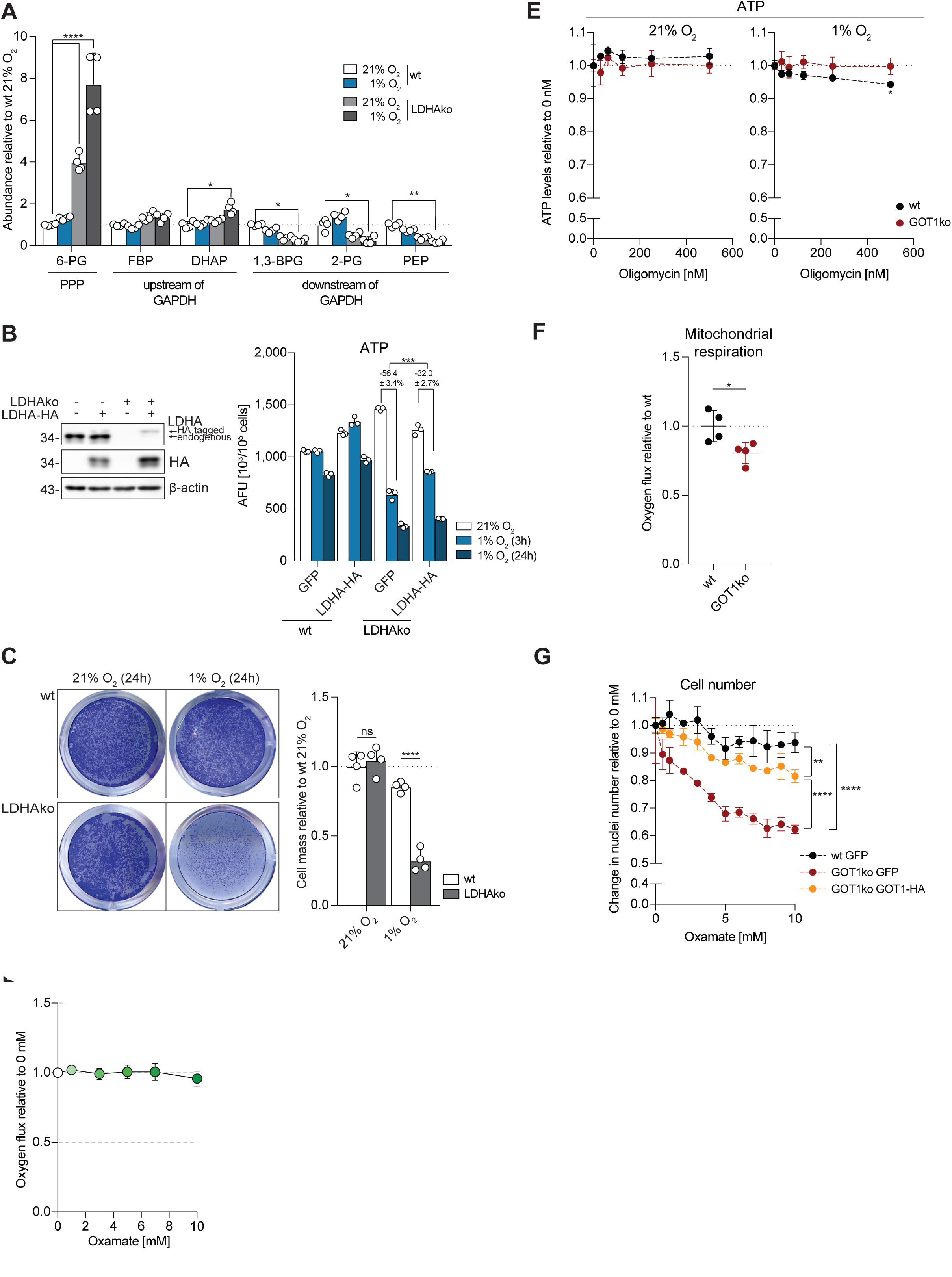
Effects of LDHA knock-out on metabolism and cell mass accumulation. **A)** Bar graph representation of the data in Figure 5C. Data shown as mean ± SD (n = 4 cultures per condition) and significance was tested using two-way ANOVA with Sidak’s multiple comparison test. **B)** Left side: Western blot to assess the levels of endogenous LDHA and HA-tagged LDHA in wild-type and LDHAko MCF7 cells stably expressing LDHA-HA or GFP. Right side: ATP levels in the same cell lines at 21% O_2_ and after 3 h or 24 h in 1% O_2_. Data shown as mean ± SD (n = 3 assays per cell line condition). Statistical errors were propagated to calculate the error of the change in ATP levels between normoxia and 3 h hypoxia and significance was tested using a two-tailed, unpaired t-test. See also Figure 5E. **C)** Cell mass accumulation in wild-type (wt) and LDHAko MCF7 cells assessed by crystal violet staining after 24 h at 21% O_2_ or 1% O_2_. Left side shows images of representative wells (with fixed and stained cells) from under each condition; right side shows quantification of crystal violet staining shown as mean ± SD (n = 3 replicates per cell line and condition) and significance was tested using two-way ANOVA with Tukey’s multiple comparison test. **D)** Cellular oxygen consumption in MCF7 cells treated successively with increasing oxamate concentrations. Data shown as mean ± SD (n = 2-3 replicates per concentration) and significance relative to 0 mM was tested using one-way ANOVA with Dunnett’s multiple comparison test (not significant). **E)** ATP levels in wild-type (wt) and GOT1ko MCF7 cells at 21% O_2_ and after 3 h in 1% O_2_ treated with the indicated oligomycin concentrations for 3 h. Data shown as mean ± SD (n = 3 assays per cell line condition) and significance relative to 0 mM was tested ANOVA with Dunnett’s multiple comparison test. **F)** Mitochondrial respiration of wild-type (wt) and GOT1ko MCF7 cells. Cellular oxygen consumption was corrected for ROX (residual oxygen consumption) by addition of the complex III inhibitor antimycin A. Data shown as mean ± SD (n = 4 replicates per condition) and significance was tested an unpaired, two-tailed t-test. **G)** Change in cell confluence of wild-type (wt) and GOT1ko MCF7 cells stably expressing GOT1-HA or GFP within 24 hours at 21% O_2_ or 1% O_2_ with the indicated oxamate concentration in cell culture media, shown relative to 0 mM oxamate per cell line. Data shown as mean ± SD (n = 2-3 cultures per cell line and condition) and significance relative to 0 mM was tested ANOVA with Tukey’s multiple comparison test (only shown for 10 mM oxamate).

**Figure S6 related to Figure 6.**
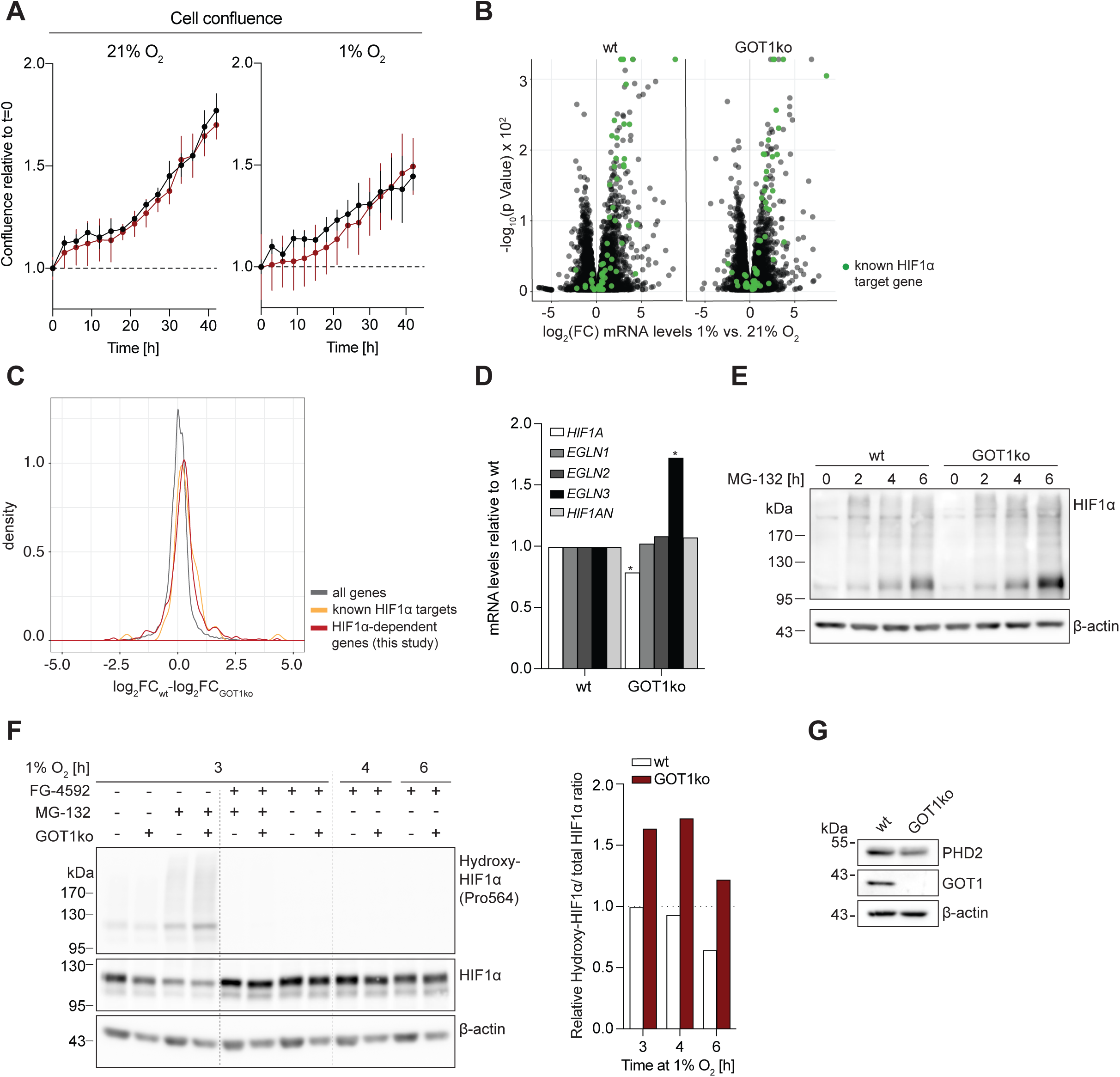
HIF1α hydroxylation, stability and target gene expression in GOT1ko cells. **A)** Cell confluence of wild-type (wt) and GOT1ko MCF7 cells at 21% O_2_ or 1% O_2_, shown relative to t=0 h. Time points also indicate duration of hypoxia treatment. Data shown as mean ± SD (n = 3 cultures per cell line and condition) and significance relative to wt cells was tested using two-way ANOVA with Sidak’s multiple comparison test (not significant). **B)** Volcano plot of gene expression changes in wild-type (wt) MCF7 and GOT1ko cells exposed to 1% O_2_ for 24 h, compared to control cells in normoxia (n = 3 cultures per condition; only changes with FDR<0.01 are shown). HIF1α target genes shown in green. **C)** Histogram showing frequency density of the difference in log_2_-fold mRNA expression changes after 24 h in 1% O_2_ between wild-type (wt) and GOT1ko MCF7. The graph shows the distribution of all genes, known HIF1α target genes and experimentally determined HIF1α-dependent genes from this study (See **Table S1**). HIF1α-dependent genes are defined as protein-coding genes (excluding long non-coding RNAs, antisense RNAs and miRNAs), the expression of which changed more than 2-fold in wild-type MCF7 cells after 24 h at 1% O_2_ but less than 2-fold in HIF1α^mut^ MCF7 cells. **D)** Fold-change of expression of the indicated genes in GOT1ko cells relative to wild-type (wt) MCF7 cells. *EGLN1*, *EGLN2*, *EGLN3* and *HIF1AN* encode PHD2, PHD1, PHD3 and factor inhibiting HIF (FIH), respectively. Asterisks indicate genes with FDR<0.01. **E)** Western blot to assess the levels of HIF1α in wild-type (wt) and GOT1ko MCF7 cells treated with the proteasome inhibitor MG-132 (10 μM) for the indicated lengths of time. **F)** Western blot to assess the levels of HIF1α and hydroxylated HIF1α (Pro564) in wild-type (wt) and GOT1ko MCF7 cells exposed to 1% O_2_ for the indicated lengths of time. Cells were treated with the PHD inhibitor FG-4592 (50 μM), the proteasome inhibitor MG-132 (10 μM) or a combination of both for the duration of the experiment. Right side shows quantification of lanes corresponding to MG-132–only treated samples from western blots in Figure 6E and **S6F**, presented as the ratio of hydroxylated HIF1α to total HIF1α. See also Figure 6E. **G)** Western blot to assess the levels of PHD2 in wild-type (wt) and GOT1ko MCF7 cells.

**Table S2:** Table of fragment ions used for metabolite quantification by GC-MS.

| Metabolite | KEGG ID | m/z | Chemical Formula | Notes |
| --- | --- | --- | --- | --- |
| 3-Phosphoglycerate (3PG) | C00197 | 459 |  |  |
| Alanine | C00041 | 116 |  |  |
| Aspartate | C00049 | 334 | C12 O4 N1 H28 Si3 |  |
| Citrate | C00158 | 375 | C15 O5 H31 Si3 |  |
| Cysteine | C00097 | 220 |  |  |
| Fumarate | C00122 | 245 | C9 O4 H17 Si2 |  |
| Glutamate | C00025 | 348 | C13 O4 N1 H30 Si3 |  |
| α-Glycerophosphate (α-GP) | C03189 | 445 | C14 O6 H38 Si4 | |
| Glycine | C00037 | 174 |  |  |
| Isoleucine | C00407 | 158 |  |  |
| Lactate | C00186 | 117 | C5 O1 H13 Si1 |  |
| Leucine | C00123 | 158 |  |  |
| Lysine | C00047 | 217 |  |  |
| Malate | C00149 | 245 | C10 O3 H21 Si2 | Used in <sup>2</sup>H-labelling experiments |
| Malate | C00149 | 335 | C12 O5 H27 Si3 | Used in <sup>13</sup>C-labelling experiments |
| Methionine | C00073 | 176 |  |  |
| *myo*-Inositol | C00137 | 318 |  |  |
| Ornithine | C00077 | 174 |  |  |
| Phenylalanine | C00079 | 218 |  |  |
| Proline | C00148 | 142 |  |  |
| Putrescine | C00134 | 174 |  |  |
| Pyruvate | C00022 | 174 | C6 O3 N1 H12 Si1 |  |
| *scyllo*-Inositol, internal standard | C06153 | 318 |  |  |
| Serine | C00065 | 204 |  |  |
| Succinate | C00042 | 247 | C9 O4 H19 Si2 |  |
| Threonine | C00188 | 320 |  |  |
| Tryptophan | C00078 | 218 |  |  |
| Tyrosine | C00082 | 218 |  |  |
| Uracil | C00106 | 241 |  |  |
| Valine | C00183 | 144 |  |  |

**Table S3:** Table of fragment ions used for metabolite quantification by LC-MS.

| Metabolite | KEGG ID | Polarity | m/z | Fragments | Chemical Formula |
| --- | --- | --- | --- | --- | --- |
| 6-Phosphogluconate | C00345 | - | 275.0173 | 96.969 | C6 O10 H13 P1 |
|  |  |  |  | 78.959 |  |
| Fructose 1,6-bisphosphate | C00354 | + | 341.0033 | 96.970 | C6 O12 H14 P2 |
|  |  |  |  | 78.959 |  |
| Dihydroxyacetone phosphate | C00111 | - | 168.9908 | 96.970 | C3 O6 H7 P1 |
|  |  |  |  | 78.959 |  |
| 1,3-Bisphosphoglycerate | C00236 | - | 264.9520 | 116.975 | C3 O10 H8 P2 |
|  |  |  |  | 78.959 |  |
| 2-Phosphoglycerate | C00631 | - | 184.9857 | 78.958 | C3 O7 H7 P1 |
|  |  |  |  | 96.969 |  |
|  |  |  |  | 166.975 |  |
| Phosphoenolpyruvate | C00074 | - | 166.9751 | 78.958 | C3 O6 H5 P1 |

