## Supplementary Table 4 for "GOT1 primes the cellular response to hypoxia by supporting glycolysis and HIF1α stabilisation"

| **Oligonucleotides** | **Source** | **Identifier** |
| --- | --- | --- |
| Forward primer GOT1 cDNA amplification: cgcacgcgtaccATGGCACCTCCGTCAGTC | This paper | N/A |
| Reverse primer GOT1 cDNA amplification: gcgctcgagCTGGATTTTGGTGACTGCTTC | This paper | N/A |
| Forward primer LDHA cDNA amplification: cgcacgcgtaccATGGCAACTCTAAAGGATCAG | This paper | N/A |
| Reverse primer LDHA cDNA amplification: gcgctcgagAAATTGCAGCTCCTTTTGGATC | This paper | N/A |
| Forward primer HIF1α CRISPR sgRNA: caccgTTCTTTACTTCGCCGAGATC | This paper | N/A |
| Reverse primer HIF1α CRISPR sgRNA: aaacGATCTCGGCGAAGTAAAGAAc | This paper | N/A |
| Forward primer GOT1 CRISPR sgRNA: caccgAGTCTTTGCCGAGGTTCCGC | This paper | N/A |
| Reverse primer GOT1 CRISPR sgRNA: aaacGCGGAACCTCGGCAAAGACTc | This paper | N/A |
| Forward primer GOT2 CRISPR sgRNA: caccgTGGAAGGCGGCGGCGATCCC | This paper | N/A |
| Reverse primer GOT2 CRISPR sgRNA: aaacGGGATCGCCGCCGCCTTCCAc | This paper | N/A |
| Forward primer LHDA CRISPR sgRNA: caccGGCTGGGGCACGTCAGCAAG | This paper | N/A |
| Reverse primer LHDA CRISPR sgRNA: aaacCTTGCTGACGTGCCCCAGCC | This paper | N/A |
| Forward primer HIF1α knockout validation: TTCCATCTCGTGTTTTTCTTGTTGT | This paper | N/A |
| Reverse primer HIF1α knockout validation: CAAAACATTGCGACCACCTTCT | This paper | N/A |
| M13 forward primer: TGTAAAACGACGGCCAGT | This paper | N/A |
